# Internalization of exogenous myelin by oligodendroglia promotes lineage progression

**DOI:** 10.1101/2025.07.18.665370

**Authors:** Carla Peiró-Moreno, Juan Carlos Chara, Katy Marshall-Phelps, Irune Ugarte-Arakistain, Stefano Calovi, Rafael Gois De Almeida, María Domercq, Carlos Matute

**Author notes:** ***Corresponding authors:*** Carlos Matute / Maria Domercq **Email:** /.

## Abstract

Oligodendrocytes, traditionally recognized for their role in central nervous system myelination, have emerged during the last decades as key participants maintaining brain homoeostasis in response to metabolic demands and stress. In addition, injury to myelin prompts a regenerative response that leads to the formation of new myelin sheaths. However, the signals regulating effective remyelination by oligodendrocytes are still not completely understood. Here, we report that oligodendrocytes can internalize exogenous myelin both *in vitro* and *in vivo*, which leads to an increase in their proliferation and differentiation when their functions are not compromised. RNA sequencing reveals that myelin debris alters oligodendrocyte transcriptional profile, suppressing immune-related pathways and *de novo* cholesterol and fatty acid biosynthesis, while inducing lipid droplet formation to store and process internalized myelin particles. As a result, progression of the oligodendroglial lineage is enhanced in primary cell cultures, as shown by increased viability, proliferation and differentiation. Oligodendrocytes also acquire a more differentiated phenotype, with larger cell areas, a more complex morphology and myelination of synthetic nanofibers. Stereotaxic injection of fluorescent myelin into mouse cortex shows internalization by microglia and, to a lesser extent, by oligodendroglia. Notably, in the zebrafish model, ventricular injections of myelin also increase the number of ventral oligodendrocytes in the spinal cord, further supporting that myelin can promote lineage progression. These findings challenge the classical view that myelin debris intrinsically inhibits oligodendrocyte proliferation, suggesting instead that oligodendrocytes can use myelin to support self-renewal and maturation, acting as a trophic factor in the absence of pathological cues.

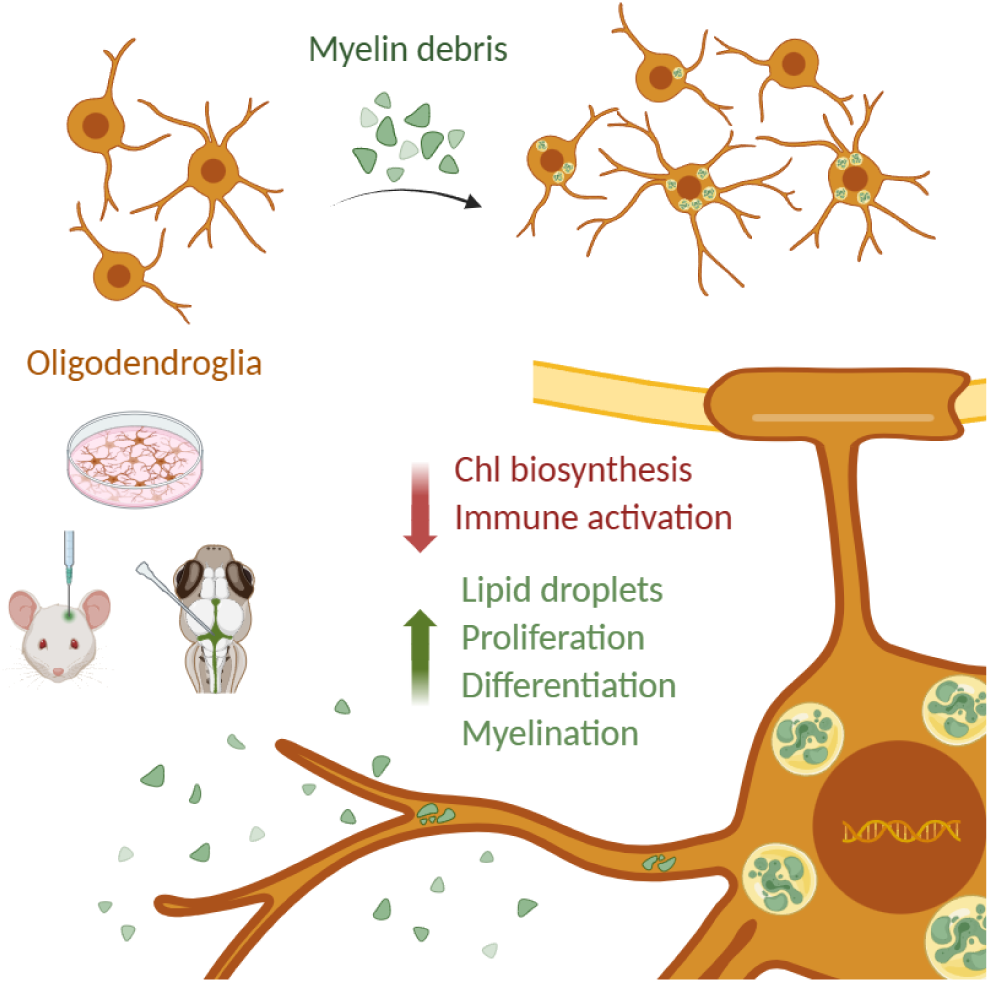

## Introduction

Oligodendrocytes (OLs) are a highly specialised class of glial cells of central nervous system (CNS), best known for their role in producing myelin sheaths following their differentiation from oligodendrocyte progenitor cells (OPCs) during early development (Bradl and Lassmann 2010; Emery 2010; Nave and Werner 2014). Myelin is a multilamellar, lipid-rich structure composed of approximately 70–80% lipids, 20% proteins and a small fraction of mRNA (Simons and Nave 2016; Stadelmann et al. 2019). Traditionally, OLs were considered primarily responsible for axonal myelination, a process that enables saltatory conduction, in which electrical impulses jump between nodes of Ranvier along the axon, allowing rapid propagation of action potentials (Bradl and Lassmann 2010; Nave and Werner 2014; Simons and Nave 2016). However, research over the last decades has expanded this view, revealing additional mechanisms through which OLs contribute to the maintenance of neuronal function.

Beyond their role in myelination, OLs also play a key role in the metabolic support of axons (Nave, Asadollahi, and Sasmita 2023; Philips and Rothstein 2017). Neurons have high energy demands due to constant ATP-dependant activity of ion pumps, synaptic vesicle recycling and generation of membrane potentials (Faria-Pereira and Morais 2022). Paradoxically, their lack of intrinsic energy reserves makes them reliant on surrounding glial cells for metabolic support. In addition to astrocytes (Tekkök et al. 2005), under homeostatic conditions OLs contribute to this energy supply primarily by transferring lactate to axons through the monocarboxylate transporter 1 (MCT1) (Lee et al. 2012; Rinholm et al. 2011). Furthermore, it has recently been suggested that in response to stress, OLs can modulate their own metabolic machinery to obtain energy from myelin-derived fatty acids. Genetically starved mice, lacking GLUT1 expression in OLs, exhibit a reduction in the thickness of their myelin sheaths (Asadollahi et al. 2024), a phenomenon that correlates with the loss of myelin in specific brain areas in humans following the extreme physical stress of running a marathon (Ramos-Cabrer et al. 2025). Likewise, the use of myelin lipids as an energy source has been proposed in the context of aging as a compensatory mechanism for age-related metabolic decline (Klosinski et al. 2015).

In experimental autoimmune encephalomyelitis, a mouse model of multiple sclerosis (MS), single-cell RNA analysis demonstrates that OLs exhibit a form of non-professional phagocytic behaviour, upregulating interferon response genes and expressing major histocompatibility complex type 1 and 2 (Falcão et al. 2018). Conventionally, clearance of myelin debris has been attributed to microglia and infiltrating macrophages, which actively engulf and degrade lipid-rich remnants during injury or disease (Franklin and Ffrench-Constant 2017; Kent and Miron 2024). It has been proposed that OL-lineage cells can internalize extracellular myelin debris, activating memory and effector CD4^+^ cells (Falcão et al. 2018). Therefore, there is emerging evidence that OLs may have alternative roles in pathological conditions.

Further research is needed to delineate the molecular mechanisms involved in this phagocytic response and the extent to which these non-canonical properties of OLs occur in physiological and pathological conditions. In this study, we demonstrate that oligodendroglia can internalize extracellular myelin debris both *in vitro* and *in vivo* and that this process promotes their proliferation and differentiation. By demonstrating the capacity of OLs to engage in debris clearance and that this process is associated with increased OL proliferation, we highlight the importance of a previously underappreciated aspect of their biology. This insight underscores the evolving view of OLs not just as passive support cells, but as dynamic regulators of CNS integrity and repair.

## Material and methods

### 1. Animals

#### 1.1. Rodents

Primary cell culture experiments were conducted using Sprague Dawley rats, sourced from the Animal Facility of the University of the Basque Country (UPV/EHU) (Leioa, Spain). C57BL/6J mice obtained from Jackson Laboratories (Bar Harbor, Maine, USA) were used to perform stereotaxic injections. Animals were housed under standard conditions, maintaining a 12 h light/dark cycle and *ad libitum* access to food and water. Experiments were reviewed and approved by the internal Animal Ethics Committee of the UPV/EHU, according to the guidelines set by the European Communities Council Directive 2010/63/EU.

#### 1.2. Zebrafish

Zebrafish (*Danio rerio*) were maintained under standard laboratory conditions at the University of Edinburgh, in compliance with UK Home Office regulations (PP0103366) and institutional ethical guidelines. Adult fish were housed on a 14-hour light/10-hour dark cycle. Embryos were raised at 28.5 °C in 90 mm Petri dish with 10 mM HEPES-buffered E3 medium (5 mM NaCl, 0.17 mM KCl, 0.33 mM CaCl₂, 0.33 mM MgSO₄), staged by days post-fertilization (dpf) according to Kimmel et al. (1995), and analyzed up to 4 dpf, prior to the onset of sexual differentiation.

The following transgenic lines were used in this study: *Tg(mbp:EGFP-CAAX)* (Almeida et al. 2011)*, Tg(mbp:EGFP)* (Almeida et al. 2011)*, Tg(mpeg1:EGFP)* (Ellett et al. 2011) and *Tg(olig1:nls-mApple)* (Marisca et al. 2020).

### 2. Oligodendrocyte cell culture

Primary oligodendrocyte cultures were obtained as previously described (Barres et al. 1992), with modifications (Sánchez-Gómez et al. 2018). Briefly, optic nerves were extracted from P11– 12 rats. After enzymatic and mechanical digestion, cells were filtered and seeded on 1 μg/ml poly-D-lysine–coated (PDL) coatings (Sigma-Aldrich). Otherwise stated, OLs were maintained in differentiation media, described in (Sánchez-Gómez et al. 2018).

Oligodendrocytes were seeded at 10,000 cells/well for immunocytochemical (ICC), cell viability experiments and on Ibidi μ-Dishes for time-lapse imaging. For RNAseq analysis, 300,000 cells/well were seeded. For myelination assays, cells were seeded on coverslips with polycaprolactone nanofibers (Sigma-Aldrich) at a 20,000 cells/well density and CNTF and NT-3 were omitted from the medium.

### 3. Cortical glial cell cultures

Primary mixed glial cultures were obtained from the cortical lobes of P0-2 rats, according to previously described by Mcarthy & De Vellis (1980) with modifications (Sánchez-Gómez et al. 2018). Briefly, forebrains were dissected to isolate the cortical lobes and after enzymatic and mechanical digestions, cell suspension was seeded in 75 cm^2^ flasks coated with PDL (Sigma-Aldrich). The resulting glial culture contained OPCs, microglia and astrocytes. After 14 days, the different cell types were isolated and seeded using different mechanical shakings, based on the different adhesion properties as described in Domercq et al. (2007) for microglial cells and in Sánchez-Gómez et al. (2018) for OPCs. After detachment and isolation of microglia and OPCs, the astrocyte monolayer was trypsinized with Trypsin-EDTA (0.5 g/l porcine trypsin and 0.2 g/l EDTA, Sigma-Aldrich) during 7 minutes and the cell suspension was seeded.

All cell types were seeded on PDL coatings and cell density was adjusted to experimental requirements. For ICC and cell viability assays, 10,000 OLs or microglia and 250,000 astrocytes were seeded per well and in Ibidi μ-Dishes for time-lapse imaging. Myelin was added at 1 DIV.

### 4. Hippocampal neuronal cultures

Hippocampal neurons were prepared from embryonic day 18 rat embryos. Hippocampi were dissected from embryonic brains and dissociated in TrypLE Express (Thermo-Fisher) for 10 min at 37°C. Cells were resuspended and homogenized in Neurobasal (Gibco) with 10% FBS Hyclone, 2 mM L-glutamine (both from Sigma-Aldrich) and 50 U/ml penicillin-streptomycin (Gibco). Hippocampal neurons were cultured on PDL-coated coverslips at 20.000 cells/well. The medium was supplemented with B27 (Gibco) and 20 μM 5-fluorodeoxyuridine and uridine (Sigma Aldrich) and changed every 3 days. Myelin was added at 7 DIV.

### 5. Myelin extraction and labelling

Myelin was isolated from adult rat or mouse brains following the protocol established by Norton & Poduslo (1973). Briefly, brain tissue was mechanically homogenized in 0.32 M sucrose and subjected to rounds of ultracentrifugation using 0.85 M sucrose gradients. Concentration of isolated myelin was determined using Bradford protein assay (ThermoFisher) and labelled with Alexa488-NHS or Alexa594-NHS dye (Invitrogen) for 1 hour at room temperature in PBS (pH=8). Excess dye was removed by 24-hour dialysis, and labeled myelin was resuspended in sterile PBS (pH=7.4) and frozen at -80°C.

For *in vitro* assays, myelin was thawed and vortexed for 30 seconds to obtain uniformly sized aggregates and added to culture medium at 5 µg/ml for immunofluorescence and cell viability experiments and 1 µg/ml for time-lapse recordings. For *in vivo* experiments, myelin was sonicated during 30 minutes before use and injected at 200 µg/ml.

### 6. Cell viability assays

Cell viability was assessed using the Calcein-AM dye (Invitrogen). Cells were incubated with 1 μM of dye for 30 minutes and fluorescence was measured with a Synergy HT fluorimeter reader (Bio-Tek Instruments). Results are expressed as the relative percentage of cellular viability with respect to control conditions.

### 7. Time-lapse imaging

Alexa-488 labelled myelin was added to the culture media 15 minutes prior to the beginning of the recording to let myelin debris to settle and avoid out-of-focus images. Bright-field and fluorescence images (488nm laser) were acquired every 10 minutes in 8-10 localized points using a BioStation IM-Q microscope (Nikon). Cells were maintained at 37°C and 5% CO_2_ during the recording.

### 8. Immunochemical analysis

For ICC analysis, primary cultures were fixed in 4 % paraformaldehyde (PFA) diluted in 0.1 M phosphate buffer (PB) for 20 min. IHC was performed in free-floating sections obtained as described in section 12.2. A standard immunofluorescence protocol was used consisting of a 1 hour incubation at room temperature (RT) in blocking solution containing 4 % normal goat serum (Vector Labs) and 0.1 % triton X-100 in PBS, except for PDGFR-α antibody for which normal donkey serum was used. Cells were incubated overnight at 4 °C with primary antibodies, followed by an hour incubation at RT with AlexaFluor-conjugated secondary antibodies and DAPI (4 μg/ml; Sigma-Aldrich) for nuclei staining. After washings, coverslips were mounted on glass slides using Dako Glycergel Mounting Medium (Agilent).

Primary antibodies used for immunohistochemistry were as follows: mouse anti-MBP (1:500; Biolegend), goat anti-PDGFR-α (1:250, R&D Systems), rabbit anti-NG2 (1:500; Abcam), mouse anti-Olig2 (1:500; Milipore), rabbit anti-Ki67 SP6 (1:250, Abcam) and guinea pig anti-Iba1 (1:300; Synaptic Systems). The corresponding Alexa Fluor secondary antibodies (ThermoFisher) were used accordingly to the host species of primary antibodies at a 1:500 dilution.

### 9. Lipid droplet staining and imaging

Fixed cells were stained with Oil Red O (BioVision), following manufacturer’s instructions: cells were incubated in 60% isopropanol for 5 minutes and then stained with Oil Red O (1.8 mg/ml in dH_2_O) for 20 minutes. After consecutive washings in dH_2_O, nuclei were stained with DAPI (4 μg/ml; Sigma-Aldrich) and coverslips were mounted. Images were acquired in a Zeiss LSM880 Airyscan microscope using a 40x objective.

### 10. Confocal imaging and analysis

Immunofluorescence imaging was conducted on a Zeiss LSM880 Airyscan or Leica TCS SP8. Imaging parameters were kept constant within each experimental set. Image processing and quantification were carried out using FIJI (ImageJ) software.

To assess oligodendrocyte morphology, we performed Sholl analysis. Individual cells were segmented as regions of interest and skeletonized. Morphological complexity was measured as the number of intersections between cell processes and concentric circle templates centered on the soma.

### 11. Bulk RNA sequencing

Total RNA from OLs cultures was isolated using NZY Total RNA Isolation kit (NZYTech), according to the manufacturer’s instructions. The quantity and quality of the RNAs were evaluated using Qubit RNA HS Assay Kit (Thermo Fisher) and Agilent RNA 6000 Nano Chips (Agilent Technologies), respectively. Sequencing libraries were prepared following “TruSeq Stranded mRNA Sample Preparation Guide (Part # 15031058 Rev. E)” using the “TruSeq® Stranded mRNA Library Prep” kit and TruSeq RNA CD Index Plate (Illumina). From 400ng of total RNA, mRNA was purified, fragmented and primed for cDNA synthesis using Illumina and ThermoFisher reagents. Then, A-tailing and adaptor ligation were performed. Finally, enrichment of libraries was achieved by PCR and quantified using Qubit dsDNA HS DNA Kit (Thermo Fisher Scientific) and visualized on an Agilent 2100 Bioanalyzer using Agilent High Sensitivity DNA kit (Agilent Technologies). Using STAR software version 2.7.10b (Dobin et al. 2012), FASTQ files were aligned to Rattus norvegicus genome data base “rn6” and reads for analysed features were assigned and counted from the processed BAM files using SubRead’s FeatureCounts version v2.0.3 (Liao, Smyth, and Shi 2014). Differential expression analysis was performed with the R library DESeq2 version 1.44.0 (Love, Huber, and Anders 2014). GSEA were performed with described gene sets using gene set permutations (n = 1000) for the assessment of significance and signal-to-noise metric for ranking genes from the Molecular Signature Database (https://www.gsea-msigdb.org/gsea/msigdb/index.jsp) (Liberzon et al. 2015).

### 12. Exogenous myelin internalization in mice

#### 12.1. Stereotaxic injections

Brain stereotaxic injections were performed in 8-10 weeks old C57BL/6J mice. Two 0.5 μl injections of mouse-derived Alexa488-labelled myelin diluted in saline solution (0.9% NaCl) were performed at the following coordinates: Bregma 0.5 mm AP, 1.0 mm LM, 1.3 mm VD and Bregma 2.5 mm AP, 0.8 mm LM, 1.0 mm VD. The content was injected at a constant rate of 0.1 μl/min using a Hamilton syringe, leaving the needle in place for an additional 3 minutes. Both injections were made in the same hemisphere. In sham-operated control mice, saline was injected using the equivalent procedures. Animals were perfused 24 or 48 hours after the injection.

#### 12.2. Tissue processing for immunohistochemistry

Mice were anesthetized and perfused with 4 % PFA in 0.1 M PB. Dissected brains were post-fixed overnight at 4 °C in the same solution. Then, coronal 40 µm-thick sections were obtained using a Microm HM 650V Microtome (ThermoFisher). Sections were selected for further staining based on the presence of fluorescent signal from myelin and/or the tissue scar from the surgery in the case of sham animals.

#### 12.3. Transmission electron microscopy

Mice were anesthetized and perfused with 4 % paraformaldehyde and 0.1% glutaraldehyde in 0.1 M PB. Coronal 50 μm-thick brain sections were obtained using a Microm HM650V microtome (Thermo Fisher).

To stain Alexa-488 labelled myelin using immunogold, slices were incubated with blocking solution (10% BSA, 0.1% sodium azide, 0.05% triton X-100 in TBS) for 60 min at RT. Subsequently, sections were incubated with rabbit IgG anti-Alexa 488 antibody (1:100, ThermoFisher) in blocking solution for 4 days at 4°C. Following several washes, sections were incubated for 2 h at RT with 1.4 nm gold-labeled goat anti-rabbit IgG (1:200; Nanoprobes Inc.) in blocking buffer. Sections were postfixed in 1% glutaraldehyde for 10 min at RT and washed in ddH20. Gold particles were silver-intensified with a HQ Silver kit (Nanoprobes) in the dark for 12 min and tissue was washed with ddH20 followed by 0.1 M PB.

The day after, sections were osmicated (1% OsO4 in 0.1 M PB) for 30 min and dehydrated in graded ethanol concentrations (50–100%) to propylene oxide and embedded in epoxy resin (Sigma-Aldrich) by immersion in decreasing concentration of propylene oxide. Tissue was then embedded in fresh resin overnight and allowed to polymerize at 60°C for 2 days.

Semithin sections (500 nm thick) were cut using a PowerTome ultramicrotome (RMC Boeckeler) and stained with 1% toluidine blue for sample orientation. Ultrathin sections (50–60 nm thick) were cut with a diamond knife (Diatome), collected on nickel mesh grids, and stained with 4% uranyl acetate for 30 min followed by 2.5% lead citrate for 10 min.

TEM images were obtained using the JEOL JEM 1400 Plus electron microscope (SGiker, UPV/EHU) at magnifications ranging from 1000x to 8000x.

### 13. Myelin internalization in zebrafish

#### 13.1. Intracerebroventricular injections

At 3 dpf, larvae were anesthetized using 600 μM Tricaine (Sigma-Aldrich) and embedded in a 2.5% agarose drop. Larvae were oriented ventrally, with the yolk sac oriented to the bottom of the drop, allowing access to the dorsal area. The needle was positioned through the thin roof plate of the hindbrain without damaging the brain tissue beneath. Each larva received two sequential 0.5 nl microinjections of Alexa-594 or -488 labelled myelin in nuclease-free water, which was also injected as a control. Following the injection, heartbeat and circulation were checked and larvae were gently released from the agarose. Injected fish were transferred to 12-well plates with E3, where they were maintained until imaging. Fish were monitored until full recovery from anaesthesia, and only those exhibiting normal swimming behaviour were included in subsequent analyses.

#### 13.2. Live imaging and quantification of zebrafish larvae

For live imaging, 3–4 dpf zebrafish larvae were anesthetized with 600 μM Tricaine (Sigma-Aldrich) and mounted laterally in 2.5% agarose as described by Vagionitis & Czopka (2018). Imaging was performed using a Zeiss LSM880 confocal microscope equipped with Airyscan detection and a Zeiss W Plan-Apochromat 20×/1.0 NA water-dipping objective or Zeiss Axio Imager Z1 equipped with an Apotome.2 unit and using a 10× objective. Larvae injected with labelled myelin were visually scored for distribution of fluorescence over the spinal cord area as high (continuous fluorescence >10 clusters), low (sparse <5 clusters), or medium with intermediate amounts of fluorescence. Animals lacking detectable fluorescence were excluded from experimental groups.

For internalization experiments, fluorescence was assessed throughout the spinal cord and brain, with specific regions of interest sampled for analysis. For figure preparation, maximum-intensity projections of z-stacks were generated and representative regions were cropped.

For cell counting, the anal pore was used as a reference point to image a consistent anterior region covering the full depth of the spinal cord. Individual oligodendrocytes were manually counted in each z-stack and classified as ventral or dorsal based on their contact with the Mauthner axon and normalized to the imaged length. A minimum of two independent injection rounds were analyzed. All zebrafish images are shown as lateral views of the spinal cord, with anterior to the left and dorsal at the top.

#### 13.3. Statistical analysis

Data are presented as mean ± standard error of mean (SEM) and n represents the number of animals, cultures or cells analyzed, as specified in figure legends. Statistical analyses were performed using GraphPad Prism 8 (GraphPad Software Inc). Comparisons between two groups were analysed using paired Student’s two-tailed t-test in *in vitro* experiments and unpaired in *in vivo* experiments. p values < 0.05 were considered statistically significant.

## Results

### 1. Oligodendroglia internalize exogenous myelin debris in vitro

In order to address the internalization capacity of oligodendroglial cells along the lineage, we first exposed different primary rat cell cultures to Alexa-labelled myelin debris for 48 hours (Figure 1B). Myelin was isolated from rats (Figure 1A) to prevent cross-species reactivity. Notably, both OLs and OPCs were found to internalize exogenous myelin (Figure 1C). Confocal imaging of OLs labelled with the membrane-linked Calcein-AM dye confirmed the internalization of myelin debris by oligodendroglia (Suppl. Figure 1).

**Figure 1.**
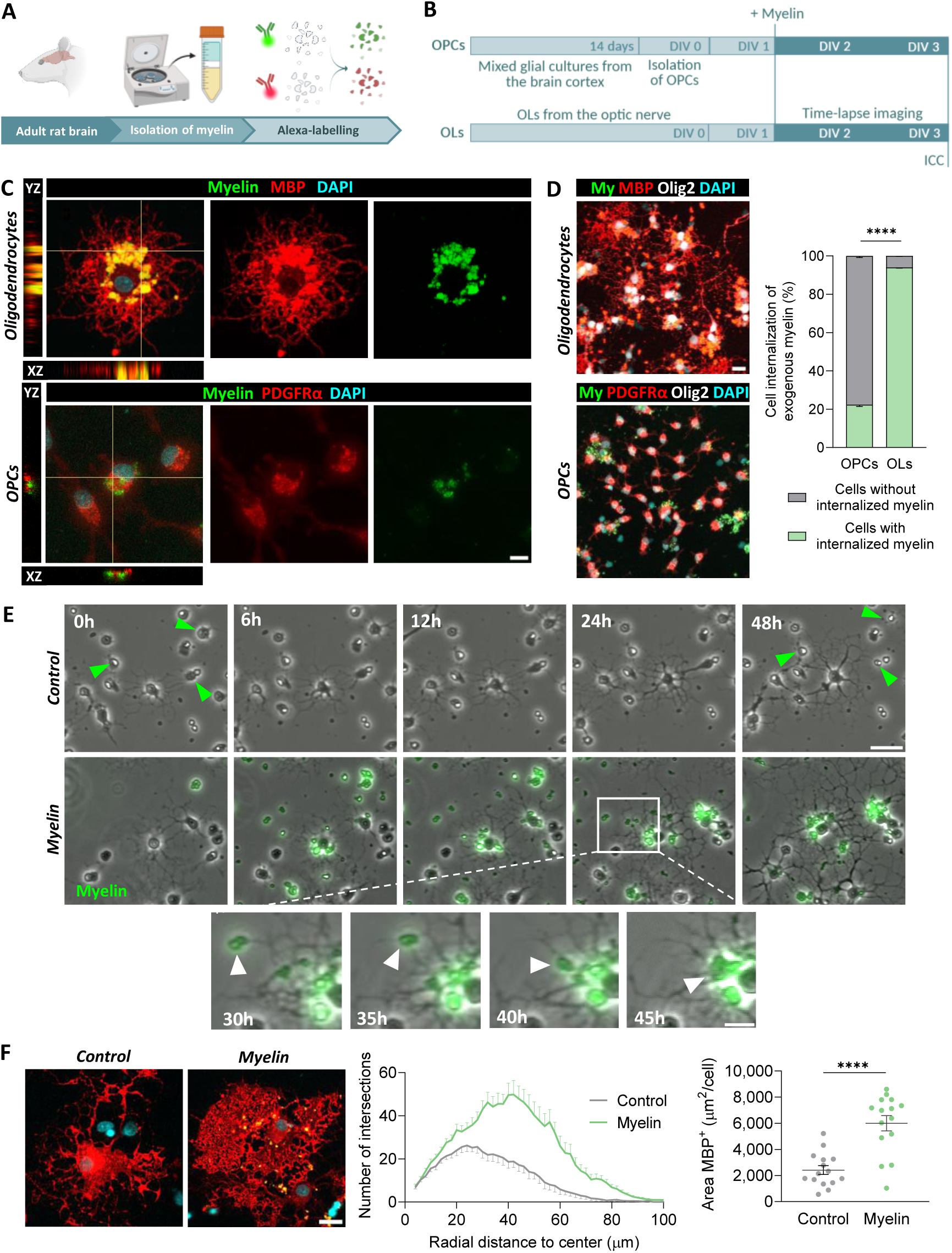
Internalization of exogenous myelin by oligodendroglial cells. (**A**) Schematic illustrating the isolation and fluorescent labelling of myelin debris. (**B**) Experimental design for evaluating myelin uptake. (**C**) Orthogonal confocal views showing internalized exogenous myelin within oligodendrocytes (OLs) and oligodendrocyte precursor cells (OPCs). Scale bar: 10 µm. (**D**) Comparative analysis of myelin internalization between OPC and OL cultures. Scale bar: 20 µm. (**E**) Time-lapse imaging of OLs reveals increased morphological complexity and cell abundance in response to myelin exposure (top), and transport of internalized myelin along OL processes (bottom). Scale bar: 50 µm; zoomed-in panel: 20 µm. (**F**) Sholl analysis of OLs after 48 hours of myelin exposure. Scale bar: 20 µm. Area under the curve: Control, 1052 ± 78.7; Myelin, 2303 ± 139; ****p < 0.0001. Peak number of intersections: Control, 26.2; Myelin, 49.9. Each dot represents a single cell (n = 15).

Interestingly, our results suggest a maturation-dependent enhancement of phagocytic or endocytic activity within the oligodendroglial lineage (Figure 1D). Specifically, OLs were more efficient at engulfing extracellular myelin, with more than 90% of cells containing internalized myelin particles. In contrast, OPCs presented significantly lower levels of internalization, with only about 20% of the cells showing uptake, potentially indicating a more limited role in debris clearance at this developmental stage.

In order to further explore the temporal dynamics of myelin internalization, we performed time lapse imaging of cultured OLs over 48 hours (Figure 1E). Interestingly, the processes of OLs were the first cellular region contacting the myelin debris and some particles were observed migrating along the processes toward the soma (Figure 1E, bottom section). The relative size of myelin debris compared to the thin OL processes strongly suggests that complete internalization occurs upon reaching the soma. After 48 hours, myelin accumulated predominantly in clusters within the cytoplasm and only small particles remained in the processes. In phagocytic cell types, such as microglia and macrophages, internalized material typically accumulates in the soma, where it is processed and degraded by lysosomal enzymes (Trivedi et al., 2020; Yu et al., 2022). Thus, the clustering of myelin within this region in OLs further supports the idea that internalization is completed upon arrival of myelin debris to the cell body.

During the 48-hour time-lapse imaging period, OLs remained static while their processes extended radially. Notably, OL processes did not exhibit directed growth toward the nearby myelin particles, suggesting that they do not actively pursue it. Instead, they appear to maintain their intrinsic growth patterns and may initiate internalization upon passive contact. Importantly, compared to controls, OLs exposed to myelin appeared more ramified and exhibited an increased cell size. Morphological sholl analysis confirmed that myelin-exposed OLs were larger and exhibited higher process complexity, as determined by increased branching. Moreover, exposure to myelin enhanced cell survival, as indicated by a reduction in cell death compared to control conditions during the time-lapse imaging period (Figure 1F, green arrowheads).

Building on these observations, we next investigated the effects of myelin exposure on microglia, astrocytes and neurons *in vitro*. As expected, microglial cells (Suppl. Figure 2A) actively phagocytosed myelin particles and over 1 hour most of microglia cell bodies visible by phase contrast progressively became Alexa-Fluor^+^. The microglial response was characterized by a higher and more rapid uptake of myelin when compared to OLs, supporting their established role in the clearance of myelin debris as professional phagocytic cells (Franklin and Ffrench-Constant 2017; Kotter et al. 2006). In contrast, astrocytes (Suppl. Figure 2B) exhibited minimal uptake and some myelin clusters were observed around the cells. Moreover, exposure to myelin did not appear to disrupt the astrocytic monolayer. Similarly, neurons showed no significant internalization of myelin (Suppl. Figure 2C). In both cases, exposure to myelin did not affect cellular survival (Suppl. Figure 2D).

These findings highlight the distinct contributions of different cell types in the internalization and clearance of myelin debris. Moreover, myelin appears to signal back specifically to OLs, promoting lineage progression and survival. These results lead us to hypothesize that myelin debris may alter metabolic and signalling pathways, thereby promoting OL differentiation and/or myelination. Our results demonstrate that OLs, in addition to microglia, are also capable of internalizing myelin, particularly as they mature. Based on these findings, we directed subsequent experiments towards myelinating OLs to further elucidate pathways triggered by myelin internalization.

### 2. Exposure to myelin alters the metabolic transcriptional profile of oligodendrocytes

We performed RNA sequencing of OLs cultured in the presence of myelin for 48 hours, which depicted 63 differentially expressed genes (Figure 2B). Gene Set Enrichment Analysis (GSEA) (Figure 2C) revealed that myelin exposure triggered a downregulation of immune-related signaling, including tumor necrosis factor alpha (TNFα), interleukin signaling, interferon response and inflammatory response and complement pathways (Figure 2D). This transcriptomic profile suggests a distinct cellular phenotype, diverging from the disease-specific state activated in OLs under experimental MS conditions (Falcão et al. 2018).

**Figure 2.**
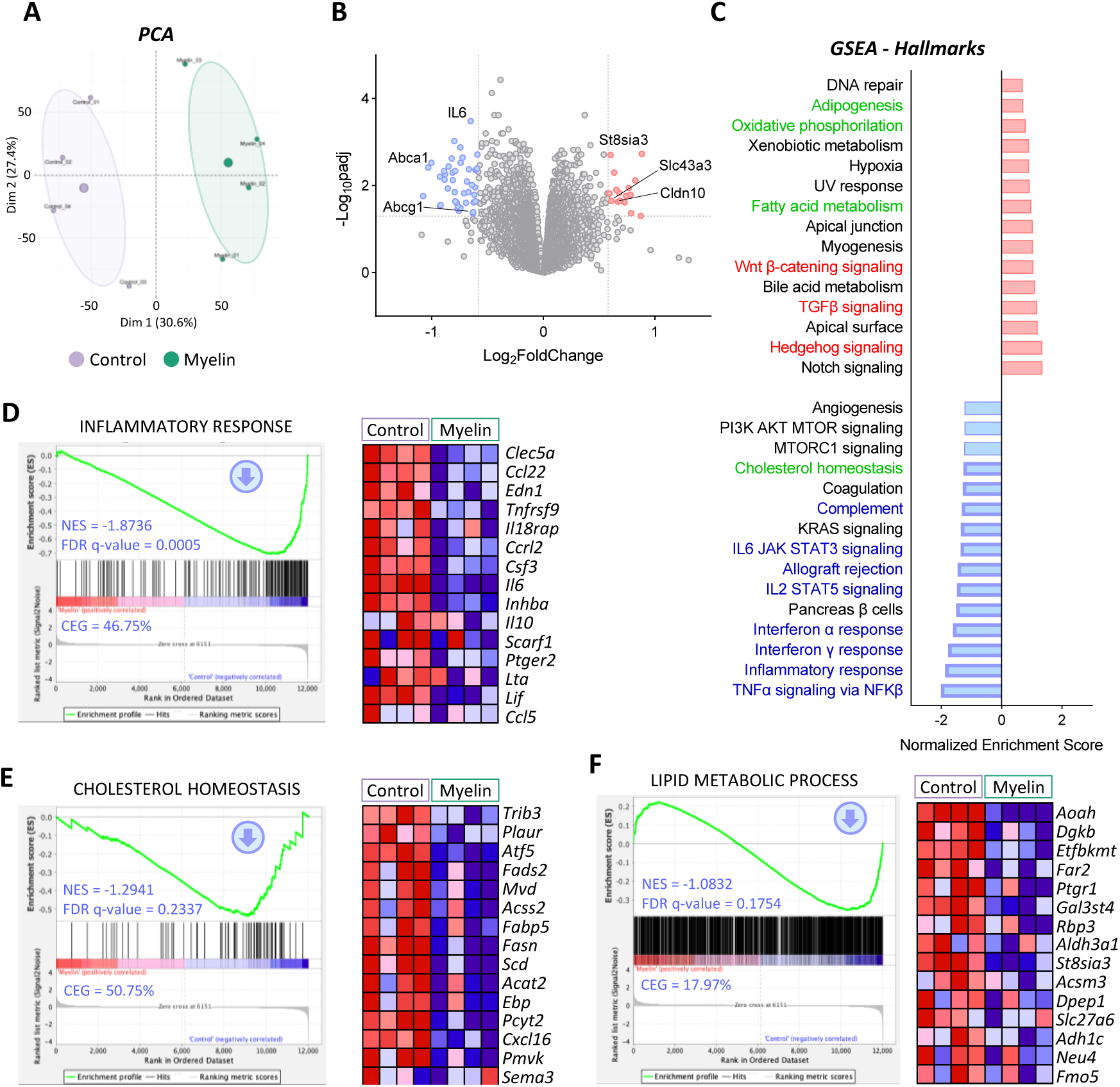
Transcriptional profile of oligodendrocytes exposed to exogenous myelin debris. OLs were treated with Alexa-labelled myelin for 48 hours, from DIV 1 to 3, followed by transcriptomic analysis (n = 4 cultures). (**A**) Principal component analysis (PCA) showing clear separation between treated and untreated samples. (**B**) Volcano plot displaying significantly upregulated genes (red) and downregulated genes (blue) in OLs exposed to myelin compared to control conditions. Differentially expressed genes (DEGs) were defined by │fold change│ > 1.5 and adjusted p-value < 0.05. Representative DEGs are labelled. (**C**) Gene Set Enrichment Analysis (GSEA) of Hallmark pathways, showing the top 15 upregulated (red) and downregulated (blue) biological processes ranked by normalized enrichment score (NES). Significantly enriched pathways (FDR q-value < 0.25) are highlighted. Metabolic, differentiation-related and immune-related pathways are indicated in green, red, and blue, respectively. (**D-E**) Enrichment plots (left) for selected Hallmark gene sets with corresponding NES, FDR q-values, and Core Enrichment Genes (CEGs), defined as the percentage of significantly enriched genes in the set. Heatmaps (right) show the top 15 most enriched genes for each pathway. (**F**) Enrichment plot for the “Lipid metabolic process” gene signature.

In addition, several metabolic pathways were significantly altered. Notably, genes involved in cholesterol homeostasis were downregulated (Figure 2E). This included reduced expression of key enzymes critical to the mevalonate pathway for cholesterol biosynthesis, such as mevalonate diphosphate decarboxylase (*Mvd*), phosphomevalonate kinase (*Pmvk*) and emopamil binding protein (*Ebp*), along with acyl-CoA synthetase short-chain family member 2 (*Acss2*), which contributes to precursor synthesis. These findings suggest that internalized myelin may serve as an exogenous source of cholesterol, thereby reducing the need for *de novo* synthesis and potentially facilitating lipid recycling for new myelin production. Indeed, Oil Red O staining revealed that internalized myelin induced an increase in lipid droplets (LDs) formation, which colocalize with myelin, suggesting that internalized myelin is stored within lipid droplets (Figure 3). LDs are key organelles that maintain lipid homeostasis, prevent cholesterol overload (Nugent et al. 2020; Ralhan et al. 2021) and regulate lipid intermediates necessary for myelin synthesis (S. A. Berghoff et al. 2021; Hayashi and Su 2004). Therefore, the presence of LDs in our model further supports the notion that internalized myelin is actively being processed and recycled.

**Figure 3.**
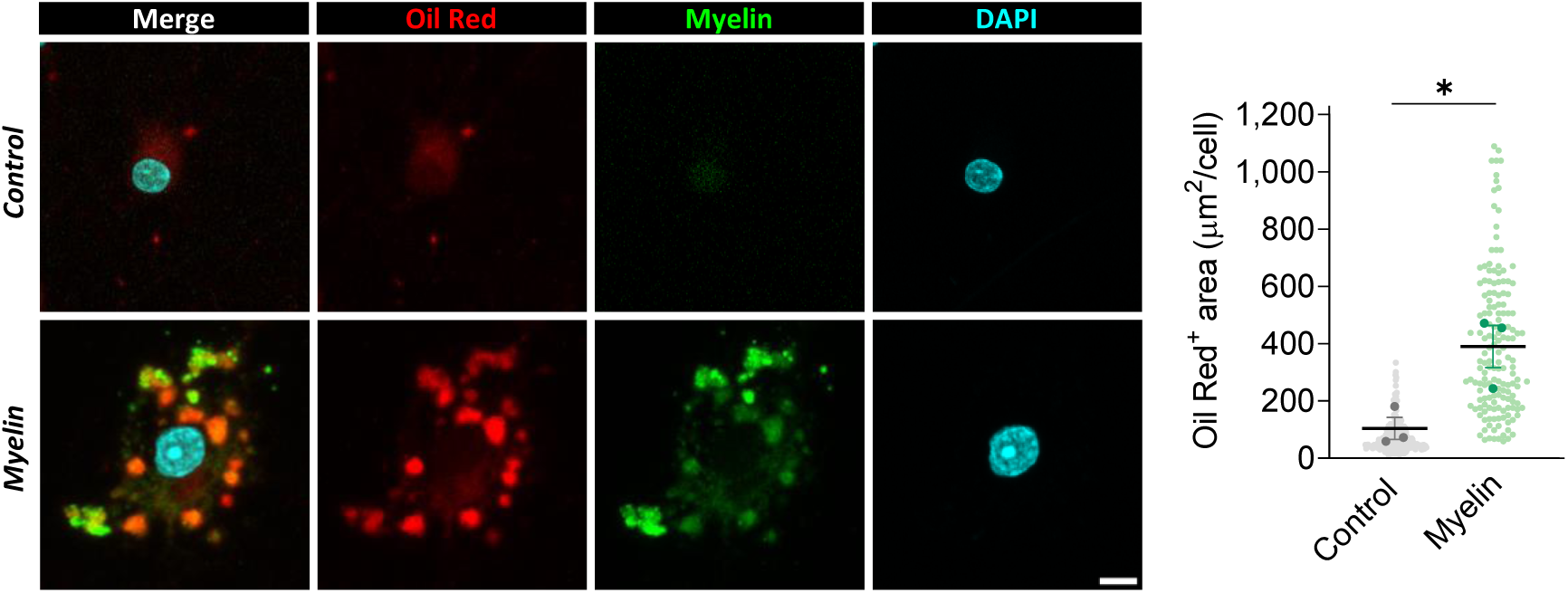
Lipid transport and storage following myelin internalization in oligodendrocytes. OLs were cultured with Alexa Fluor 488-labeled myelin for 48 hours. Myelin internalization led to increased formation of lipid droplets, visualized by Oil Red O staining, which labels neutral lipids. Scale bar: 10 µm. *p < 0.05.

Within the “Cholesterol Homeostasis” gene signature, we also found a downregulation of genes involved in fatty acid metabolism. This included fatty acid synthase (*Fasn*), a key enzyme involved in *de novo* fatty acid synthesis, along with stearoyl-CoA desaturase (*Scd*) and fatty acid desaturase 2 (*Fads2*), which mediate fatty acid desaturation and elongation, respectively. Additionally, genes linked to lipid anabolism within the “Lipid Metabolic Process” signature, such as fatty acyl-CoA reductase 2 (*Far2*) and fatty acid transporter 6 (*Slc27a6*), were also downregulated (Figure 2F). These findings suggest that myelin-derived metabolites may influence the lipid pool in OLs, thereby downregulating endogenous lipid biosynthesis pathways.

In contrast, pathways associated with lipid catabolism showed a tendency towards an enrichment, including fatty acid metabolism and oxidative phosphorylation (Figure Suppl. 3A-B). This result supports the hypothesis that OLs may be activating the molecular machinery required to process the lipid components present in internalized myelin. Among the top 15 upregulated genes within the “Fatty Acid Metabolism” signature, we identified key lipid transporters and activators, such as carnitine palmitoyltransferase 1A (*Cpt1a*) and 2 (*Cpt2*) and acyl-CoA synthetase medium-chain family member 3 (*Acsm3*). Notably, critical enzymes involved in the β-oxidation pathway, including 3-ketoacyl-CoA thiolase (*Acaa2*), 2,4-dienoyl-CoA reductase 1 (*Decr1*) and acyl-CoA thioesterase 2 (*Acot2*) and 8 (*Acot8*) were also found to be upregulated.

Finally, pathways associated with cell proliferation and differentiation, such as transforming growth factor β (*Tgfβ*) signaling, showed a non-significant enriched expression, as well as, the “Oligodendrocyte differentiation” signature (Figure Suppl. 3C-D). Altogether, transcriptomic analysis revealed that myelin exposure upregulates metabolic and signaling pathways, which could promote OL differentiation and myelination.

### 3. Myelin internalization triggers oligodendrocyte proliferation in vitro

We next assessed the effects of myelin internalization on oligodendroglial lineage progression. After exposing primary OL cultures to myelin, we found that myelin debris increased cell viability by ∼80%, as assessed by Calcein-AM, which could be attributed either to a greater number of viable cells and/or to an increase in cell size (Figure 4B). To better understand the underlying cause, we conducted additional analyses using immunocytochemistry. Consistent with cell viability assays, immunostaining of the oligodendroglial lineage marker Olig2 confirmed an increase in the total number of cells (Figure 4C). Interestingly, proliferation marker Ki67 was enhanced in cultures exposed to myelin (Figure 4C), suggesting a boost in proliferation. This observation likely reflects an expansion of the OPC pool (Dimou and Götz 2014; Young et al. 2013), which was further confirmed by the increase in NG2^+^ cells (Figure 4D). In parallel, we detected higher numbers of MBP^+^ myelinating oligodendrocytes, while the ratio of NG2^+^ to MBP^+^ cells remained constant (Figure 4D). Thus, myelin internalization accelerates lineage progression maintaining the dynamics between precursor and mature stages. These changes prompted us to investigate whether internalized myelin boosted myelination capacity of mature OLs, typically associated to a more differentiated state (Bradl and Lassmann 2010). To this end, we seeded mature OLs onto coverslips containing synthetic nanofibers. We found that exposure to myelin significantly enhanced nanofiber myelination, as shown by an increase in OL area (Figure 4E).

**Figure 4.**
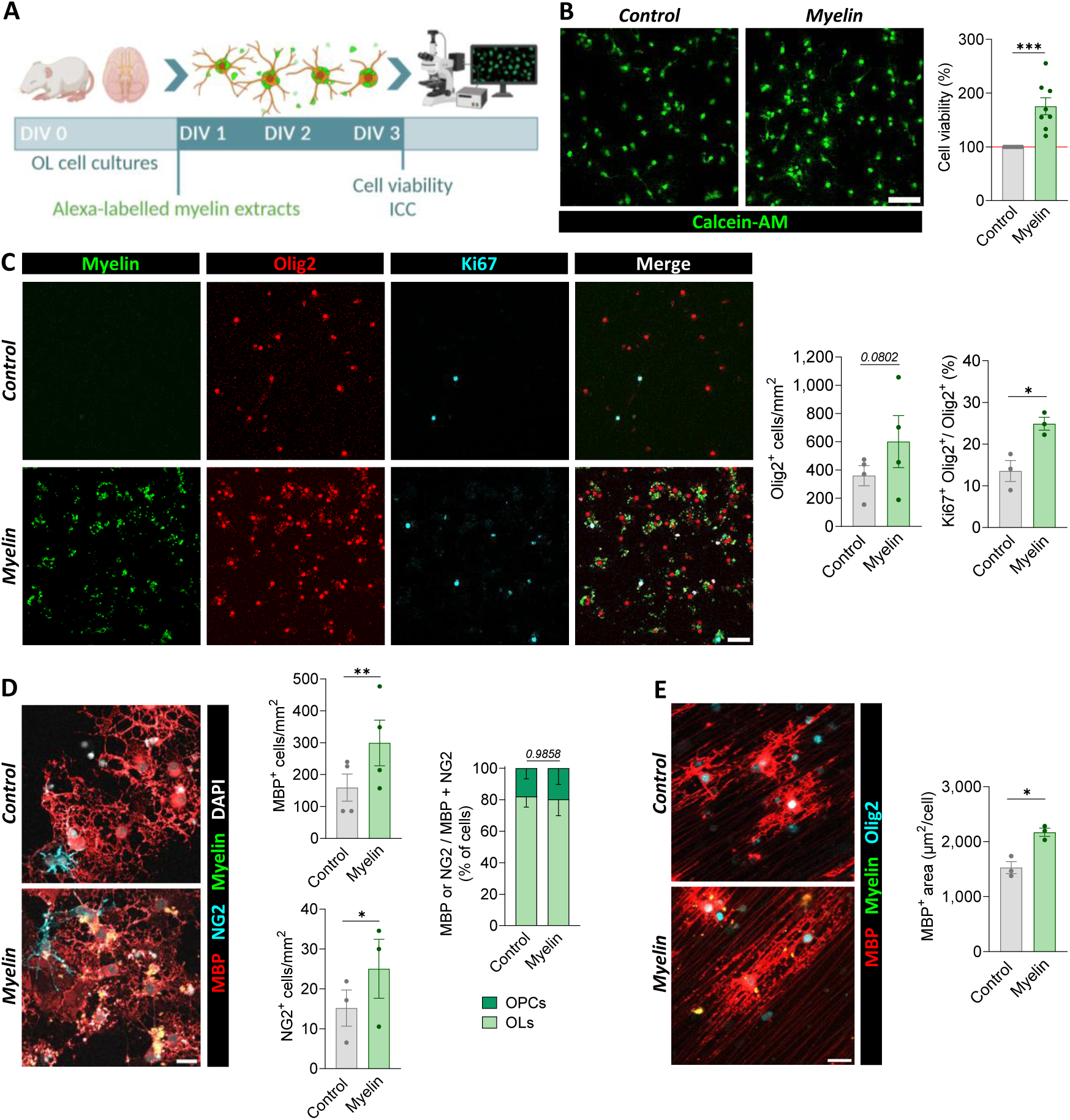
Exogenous myelin promotes oligodendroglial lineage progression. (**A**) Schematic of the experimental design. (**B**) Calcein-AM viability assay showing fluorometric values normalized to control mean. (**C**) Immunocytochemical (ICC) analysis of OL cultures indicates that myelin exposure increases the number of oligodendrocytes (Olig2⁺), associated with enhanced proliferation (Ki67⁺). (**D**) This increase results in higher numbers of myelinating oligodendrocytes (MBP⁺) and oligodendrocyte precursor cells (NG2⁺), without affecting the proportion of progenitors to mature OLs. (**E**) Enhanced myelination of nanofibers by myelin-exposed OLs, reflected by an increased cellular area. Scale bars: A–C 50 µm; D-E 25 µm. *p < 0.05, **p < 0.01, ***p < 0.001

Taken together, our results suggest that myelin internalization triggers OL proliferation, as well as morphological complexity and differentiation. Thus, myelin debris may serve as a positive regulator that activates the intrinsic program of OL development and myelination, promoting both expansion and progression of the oligodendroglial lineage.

### 4. Internalization of exogenously injected myelin in the mouse brain

*In vitro* experiments were conducted in the absence of other cell types and their associated signalling interactions present *in vivo*. Therefore, it was important to investigate whether the internalization of myelin by OLs and the accompanying increase in proliferation was also observed in a more complex cellular environment. We aimed to determine first, whether the capacity of OLs to internalize myelin is sufficiently robust to be observed in the presence of active microglia, and second, whether microglial uptake of myelin debris triggers signalling cascades that modulate OL behaviour. To this end, we performed stereotaxic injections of fluorescently labelled myelin into the cerebral cortex of adult mice. Although myelin debris is typically associated with demyelinating pathologies or injury, we sought to isolate the internalization process from confounding factors, such as OL dysfunction. While the stereotaxic procedure inherently induces a local inflammatory response, this approach enabled us to evaluate OL internalization capacity under non-demyelinating conditions. Moreover, myelin turnover may also involve clearance mechanisms beyond microglial phagocytosis, potentially implicating OLs themselves.

As expected, microglial cells rapidly migrated towards the injection site. Within the first 24 hours post-injection, most microglial cells accumulated in the injection core (Figure 5A), temporarily depleting microglia from the immediate surroundings and forming a circular gap with a radius of 150–300 µm, which was subsequently repopulated over the following 24 hours (Figure 5B). Immunohistochemistry confirmed that microglia were the predominant cell type within the injection site and internalized debris was clearly localized within phagocytic pouches (Figure 5C). Microglia adopted a rounded, amoeboid morphology typical of activated, phagocytic cells. These findings were corroborated by electron microscopy (EM), which visualized Alexa-labelled myelin debris within intracellular compartments in microglial cytoplasm (Figure 5D). This expected behaviour validated the approach and served as a positive control that the surgery procedure did not impair the capacity of phagocytic cells to internalize exogenous myelin.

**Figure 5.**
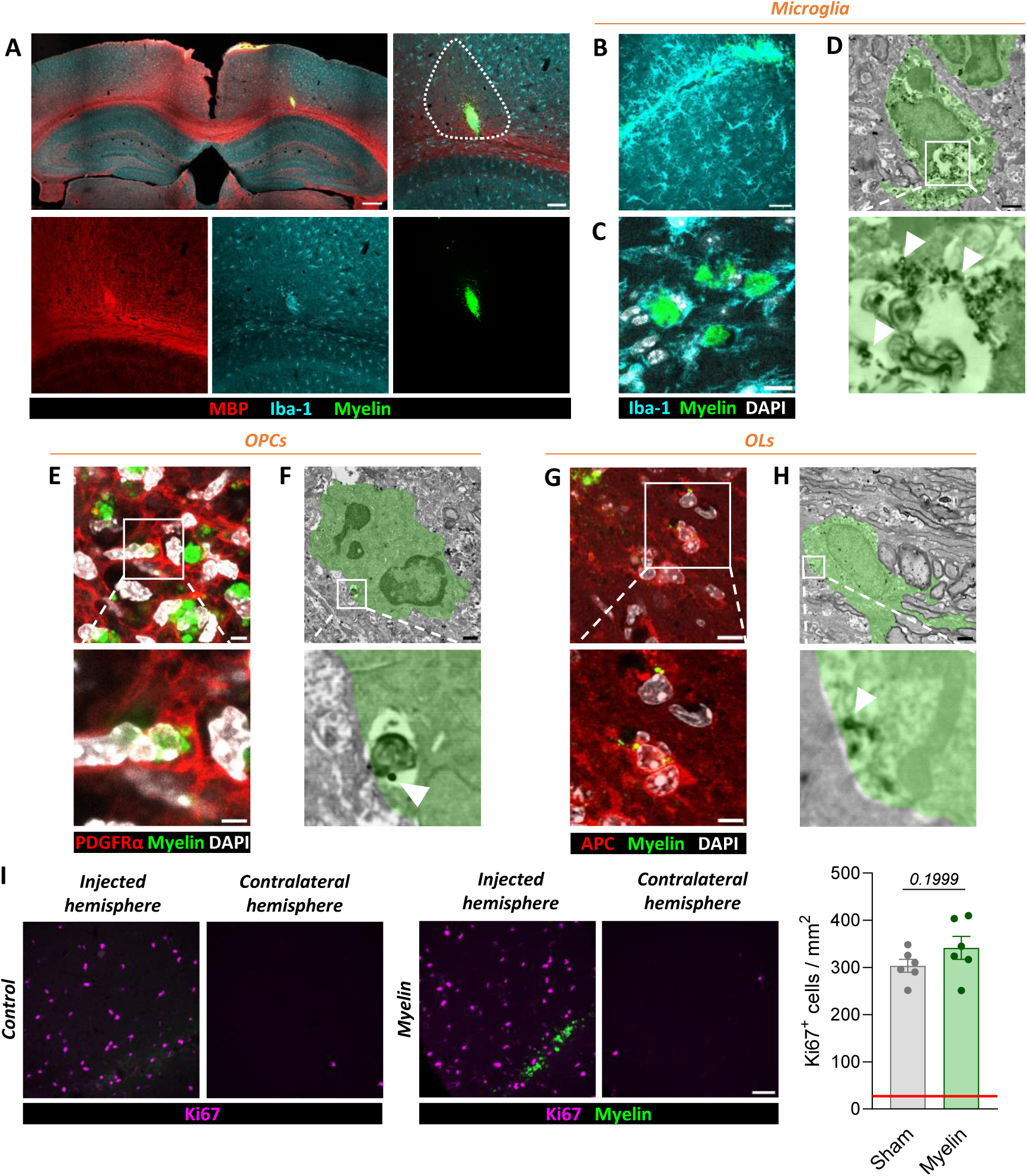
Internalization of Alexa Fluor 488-labeled myelin debris in adult mice following stereotaxic injection. (**A**) Cortical section showing microglial migration to the injection site 24 hours post-surgery, resulting in a cell-free gap around the injection core indicated by a dotted line. Scale bars: 300 µm (main), 100 µm (zoom). (**B**) Repopulation of surrounding areas by microglia 48 hours after injection. Scale bar: 50 µm. (**C**) Formation of phagocytic pouches within microglia actively internalizing myelin debris. Scale bar: 10 µm. (**D,F,H**) Immunogold labelling of Alexa Fluor 488-labeled myelin (white arrows) confirms internalization by microglia (**D**), OPCs (**F**), and oligodendrocytes (**H**), as observed by transmission electron microscopy (TEM). Microglia exhibit large vesicles (white), indicative of professional phagocytosis, whereas oligodendroglia show more diffuse internalized structures. Scale bar: 1 µm. (**E,G**) Internalization of exogenous myelin by OPCs (**E**) and OLs (**G**) *in vivo*. Scale bars: 20 µm (main), 10 µm (zoom). (**I**) Quantification of proliferating cells at the injection site 48 hours after injection in both sham- and myelin-injected animals in a radius of 300 µm around the lesion site. A dashed line indicates basal proliferation levels in a corresponding area of the contralateral hemisphere. No significant differences in basal proliferation were detected. Each dot represents an individual injection.

Interestingly, 48 hours post-injection we also identified a subset of PDGRα^+^ OPCs (Figure 5E) and APC^+^ OLs (Figure 5G) containing myelin particles, as confirmed by TEM analysis (Figure 5F and 5H). This was observed in a more limited and structurally distinct manner compared to microglia. Unlike the well-formed phagocytic vesicles seen in microglia, internalized myelin within OLs appeared in smaller, scattered inclusions without the formation of large degradation compartments. This could also reflect the lower myelin load within OLs, which may allow them to degrade and process debris without activating the same specialized degradation pathways observed in professional phagocytes.

We next sought to determine OL proliferation. The stereotaxic procedure itself triggers a local proliferative response in the cortical area surrounding the injection, masking any additional effect that myelin may exert. As a result, the proliferation marker Ki67 increased compared to the contralateral hemisphere, but no significant difference was detected between myelin- and sham-injected animals (Figure 5I). Therefore, in order to address whether myelin internalization influences OLs proliferation *in vivo* without injury-related confounds, we complemented our analysis using the zebrafish model.

### 5. Exogenously injected myelin increases oligodendrocyte number in the zebrafish

Zebrafish larvae offer a transparent and genetically tractable system allowing for high spatial-temporal resolution for live imaging (Chia et al. 2022; Doszyn, Dulski, and Zmorzynska 2024). This model enables for *in vivo* visualization of myelin internalization dynamics, while preserving the complexity of a living organism.

We performed brain ventricle injections in transgenic zebrafish larvae expressing membrane-targeted EGFP under the myelin basic protein promoter (*Tg(mbp:EGFP-CAAX*)), which allows selective labelling of OLs (Almeida et al. 2011). Membrane localization of EGFP facilitates the assessment of potential internalization of myelin, as engulfed debris can be clearly distinguished within the boundaries of myelinating oligodendrocytes, including fine processes and soma. Labelled myelin was injected into the brain ventricle at 3 days post-fertilization (dpf) and to avoid a potential confounding localized response to the injection, the spinal cord was imaged, rather than the injection site at different time points (Figure 6A). At this developmental stage, brain ventricles are already connected to the spinal canal, allowing the movement of injected material directly into the spinal cord. Control animals were injected with vehicle solution.

**Figure 6.**
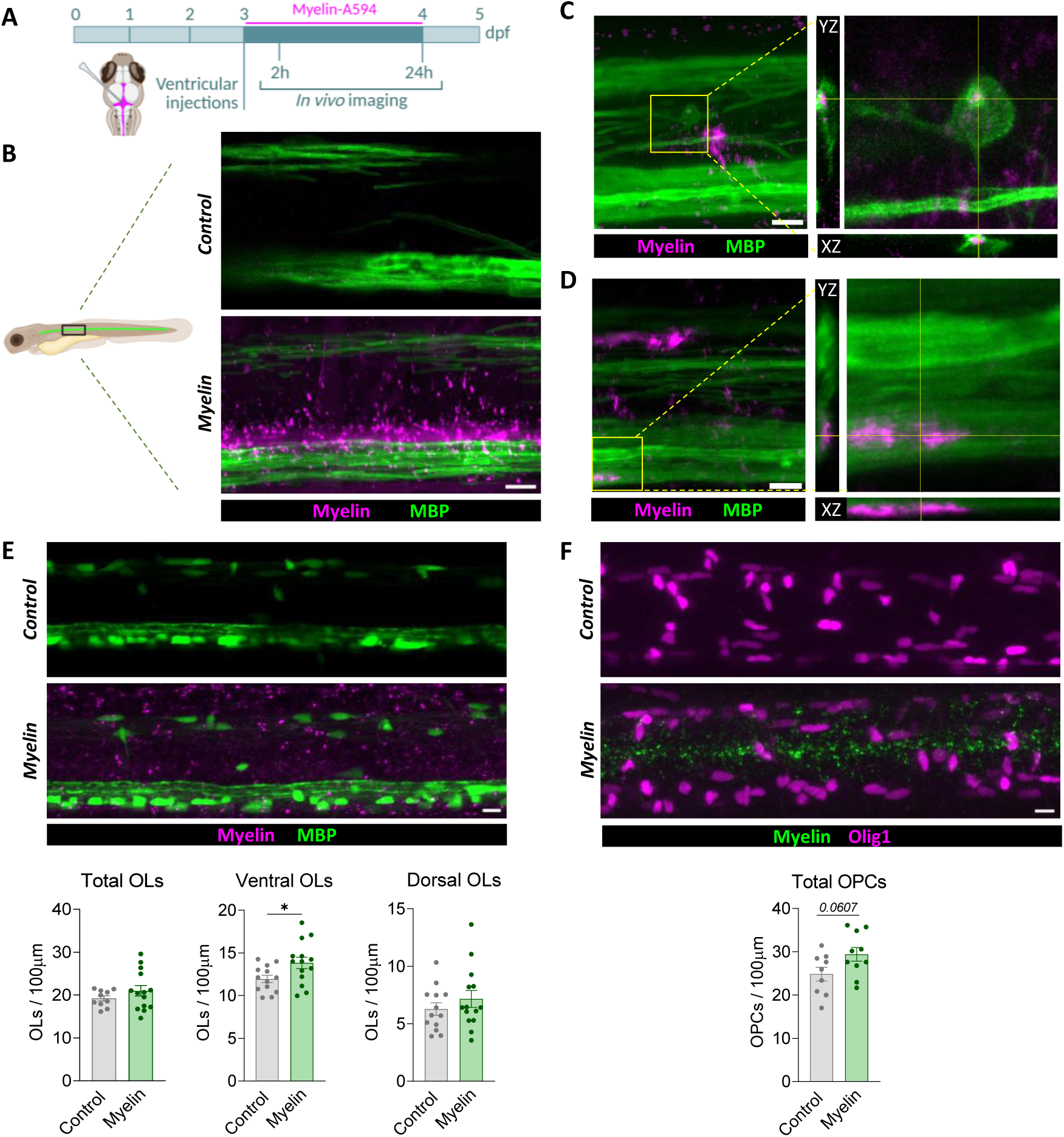
Oligodendrocytes internalize exogenous myelin in zebrafish. (**A**) Schematic representation of cerebroventricular injections of Alexa Fluor-labelled myelin performed at 3 days post-fertilization (dpf) in *Tg(mbp:EGFP-CAAX)* zebrafish, which express membrane-bound GFP in oligodendrocytes. (**B**) Confocal imaging of the spinal cord 2 hours post-injection shows the diffusion of fluorescent myelin into the central canal in the anterior region. (**C–D**) 24 hours post-injection, myelin signal is largely cleared from the canal, but a small fraction of exogenous myelin remains visible within the somata (**C**) and processes (**D**) of oligodendrocytes. (**E**) Representative spinal cord images from *Tg(mbp:EGFP)* (somatic fluorescence in OLs) and *Tg(Olig1:nls-mApple)* (nuclear marker for OPCs) zebrafish at 24 h post-injection, with quantification of myelin-positive cells. Scale bar = 10 µm.

To determine the distribution and diffusion of Alexa-labelled myelin following injection, we first imaged 2 hours post-injection. The spinal cord was divided into anterior, central and posterior regions for analysis. The anterior region (Figure 6B), located closer to the brain ventricle, showed clear presence of labelled myelin debris dispersed along and outside the central canal, reaching both the ventral and dorsal myelinated tracts of the spinal cord. The central and posterior regions (Figure Suppl 4A), situated more caudally along the tail, showed little to no detectable myelin at this time point. This confirmed that the injected material diffused through the cerebrospinal fluid and reached the anterior spinal cord, allowing for potential interaction with resident oligodendrocytes. At this early time point, however, we did not observe internalization of myelin by oligodendrocytes. Importantly, zebrafish larvae did not exhibit any visible behavioural abnormalities or morphological defects following injection and the introduced myelin was well tolerated, indicating that it was not toxic at the administered concentration.

By 24 hours post-injection, myelin fragments had formed more discrete clusters, typically 7 to 10 per fish, distributed along the spinal cord. To determine the identity of the cells containing these clusters, we repeated the injections in *Tg(mpeg1:EGFP)* zebrafish (Ellett et al. 2011), labelling microglia and macrophages. Colocalization analysis revealed that large myelin clusters were predominantly found within microglial cells, confirming their active role in myelin phagocytosis (Figure Suppl 4B). In addition to microglial uptake, we also detected smaller myelin particles contained within the soma of OLs (Figure 6C) and associated with their myelin tracts (Figure 6D). These findings demonstrate that, consistent with our *in vitro* data and murine model, OLs in zebrafish are capable of internalizing exogenous myelin.

We next evaluated the impact of myelin internalization on the oligodendroglial lineage in the zebrafish spinal cord. Remarkably, 24 hours after the injection of myelin, we observed a modest yet significant increase in the number of ventral OLs (Figure 6E), which are located closer to the central canal where the injected myelin accumulates. In contrast, dorsal OLs, potentially less exposed to myelin, did not show a significant change in cell number. This spatially restricted response suggests a direct, contact-dependent effect, potentially driven by direct internalization of myelin debris. Moreover, OPCs, imaged in *Tg(olig1:nlsmApple)* zebrafish (Marisca et al. 2020), showed a non-significant increase following injection of myelin (Figure 6F). Taken together, and considering that zebrafish development is characterized by low levels of oligodendroglial cell death (Almeida and Lyons 2016), these findings suggest that myelin uptake could locally be enhancing proliferation, rather than promoting oligodendroglial survival.

## Discussion

In this study, we found that OLs have the capacity to internalize myelin debris, leading to transcriptional changes and promoting cellular proliferation and differentiation. Traditionally, the role of myelin debris in regulating oligodendroglial dynamics has been investigated in the context of demyelination, where it is known to inhibit OPC proliferation and hinder their recruitment to lesion sites (Kotter et al. 2006; Plemel et al. 2013). However, the molecular mechanisms underlying this inhibitory effect remain incompletely understood (Baer et al. 2009; Ji et al. 2024; Molina-Gonzalez et al. 2023). Moreover, demyelination involves massive tissue damage, microglia activation and immune infiltration. Thus, signaling from damaged tissue and immune cells could potentially affect the OL population. Our findings support the idea that the deleterious impact of myelin debris is highly context-dependent, shaped by the health of OLs and signals from surrounding cellular environment. In contrast, in the absence of overt pathology, myelin may function as a trophic factor that enhances OL lineage progression.

Despite some insights, the extent to which OLs can internalize myelin and the underlying mechanisms remain insufficiently understood. Evidence suggests that OPCs may internalize myelin particles via phagocytosis (Falcão et al. 2018). Similarly, Schwann cells, the myelinating cells of the peripheral nervous system, have been shown to internalize myelin following nerve injury through specific autophagy pathways (Gomez-Sanchez et al., 2015). In the case of myelinating OLs, several studies have demonstrated that, under pathological and physiological conditions, they express several molecular components known to be involved in endocytic and phagocytic pathways. These include receptors such as low-density lipoprotein receptor-related protein 1 (LRP1), implicated in myelin uptake (Fernández-Castañeda et al. 2020; Gaultier et al. 2009; Lin et al. 2017); antigen presentation molecules (Falcão et al. 2018; Zeis, Enz, and Schaeren-Wiemers 2015); and members of the tetraspanin family (Terada et al. 2002). In this study, we demonstrated that OLs not only have the capacity to internalize myelin debris but also to transport it retrogradely from their processes to the soma, where internalization is completed and degradation begins. This observation suggest the presence of a surface-based trafficking mechanism along their processes. Consistent with this, microvesicles interacting with microglia are known to exhibit directional motion along the external plasma membrane toward the soma, a process dependent on phosphatidylserine-mediated recognition (Prada et al. 2016).

Experimental models of demyelination often involve direct targeting and damage to OLs (Blakemore and Franklin 2008; Lassmann and Bradl 2017; Torre-Fuentes et al. 2020; Zirngibl et al. 2021), which inherently disrupts their function and may obscure broader roles. In response to injury, OPCs become activated, undergoing transcriptomic reprogramming to an immature state that promotes proliferation and migration by upregulation of immune-related genes (Moyon et al. 2015). OLs also upregulate inflammatory mediators, stress response genes, anti-oxidant enzymes, cholesterol metabolism and growth factors (Hou et al. 2023). This signature largely overlaps with the disease-associated OLs recently reported in different neurodegenerative models (Pandey et al. 2022). Importantly, TREM2 deficiency in microglia impaired transcriptomic changes in disease associated OLs, suggesting that OL reprogramming is driven by microglial activation rather than by exposure to myelin debris (Hou et al. 2023). Our data shows that exposure to myelin debris, in the absence of an inflammatory insult, induces i) an opposite transcriptional signature in OLs characterized by a reduced expression of immune-related pathways and cholesterol biosynthesis and ii) an increase in OLs proliferation and differentiation. It is therefore possible that during demyelination pathological cues directly or indirectly influence OL behavior.

During CNS development, reduced expression of the cholesterol transporters Abca1 and Abcg1, transporters of cholesterol efflux to the extracellular space, is associated with the onset of active myelination (Nelissen et al. 2011). Transcriptomic analysis also suggested predominant upregulation of the cholesterol-synthesis pathway in OLs during remyelination (Voskuhl, R.R et al., 2019; Berghoff, S.A et al., 2022). This downregulation facilitates cholesterol retention within the cell, essential for myelin synthesis. A similar regulatory pattern emerges under demyelinating conditions, wherein neurons upregulate cholesterol synthesis, while OLs downregulate *Abca1* and *Abcg1* expression to minimize cholesterol efflux and prioritize lipid availability for remyelination (S. Berghoff et al. 2021; Meffre et al. 2015). Consistent with these observations, our findings demonstrate that when OLs are exposed to myelin debris, they adopt a lipid-conserving phenotype that promotes cholesterol retention and supports myelin synthesis. Similarly, lipid droplets, that regulate lipid intermediates necessary for myelin synthesis during development (Hayashi and Su 2007), are increased after myelin exposure. This adaptive response mirrors the molecular mechanisms observed during developmental myelination, suggesting a conserved strategy for optimizing lipid utilization.

While this study did not aim to dissect the molecular mechanisms through which myelin promotes OL proliferation and differentiation, this remains a key area for future research. Based on its lipid-rich composition and previous findings (Asadollahi et al. 2024; Ramos-Cabrer et al. 2025), OLs could be using myelin to buffer energetic stress. Although β-oxidation capacity of OLs is considered limited (Fünfschilling et al. 2012; Rao et al. 2017; Tepavčevic 2021), there is evidence that suggests it is still functionally active (Hofmann et al. 2017). Importantly, in contrast to previous studies, our experimental paradigm did not restrict glucose availability. Therefore, the observed effects are unlikely to be driven by a metabolic shift away from their preferentially glycolytic activity, but rather as a response to excess lipid availability. Additionally, both lipids and proteins in myelin are known to exert signaling functions. Lipid receptors such as CPT1A (Tang et al. 2022), sphingosine-1-phosphate receptor (S1PR) (Jaillard et al. 2005) or Lysophosphatidic acid receptor 1 (LPAR1) (Nogaroli et al. 2009; Yung et al. 2015) play essential roles in regulating lipid homeostasis and myelination. These pathways may be activated upon myelin internalization and could contribute to the observed proliferative and morphological responses in OLs.

In conclusion, our findings identify that OLs can internalize and respond to myelin debris, suggesting that myelin debris itself is not intrinsically hindering the progression of the oligodendroglial lineage. Rather, detrimental effects associated with myelin debris may stem from concomitant OL damage. These results underscore the critical role of OL integrity during pathology.

## Acknowledgments

This work was supported by Spanish Ministry of Education and Science (PID2019-109724RB-I00, PID2022-143020OB-I00 and PID2022-138276OB-I00), Basque Government (IT1551-22) and CIBERNED (CB06/05/00 76). C.P.M. holds a predoctoral fellowship from the Spanish Ministry of Education and Science. The authors thank to SGIker (UPV/EHU) and Achucarro facilities. K.L.H.M-P. and R.G.A. acknowledge funding support from a Chancellor’s Fellowship from the University of Edinburgh to R.G.A.; the Biotechnology and Biological Sciences Research Council [BB/X009394/1] and the UK Research and Innovation under the UK government’s Horizon Europe funding guarantee [EP/Y024311/1].

## References

1. Almeida, Rafael G., Tim Czopka, Charles Ffrench-Constant, and David A. Lyons. 2011. “Individual Axons Regulate the Myelinating Potential of Single Oligodendrocytes in Vivo.” *Development (Cambridge*, England) 138(20):4443–50. doi:10.1242/DEV.071001.

2. Almeida, Rafael, and David Lyons. 2016. “Oligodendrocyte Development in the Absence of Their Target Axons In Vivo.” PLoS ONE 11(10):e0164432. doi:10.1371/JOURNAL.PONE.0164432.

3. Asadollahi, Ebrahim, Andrea Trevisiol, Aiman S. Saab, Zoe J. Looser, Payam Dibaj, Reyhane Ebrahimi, Kathrin Kusch, Torben Ruhwedel, Wiebke Möbius, Olaf Jahn, Jun Yup Lee, Anthony S. Don, Michelle-Amirah Khalil, Karsten Hiller, Myriam Baes, Bruno Weber, E. Dale Abel, Andrea Ballabio, Brian Popko, Celia M. Kassmann, Hannelore Ehrenreich, Johannes Hirrlinger, and Klaus-Armin Nave. 2024. “Oligodendroglial Fatty Acid Metabolism as a Central Nervous System Energy Reserve.” Nature Neuroscience 27:1934–44. doi:10.1038/s41593-024-01749-6.

4. Baer, Alexandra S., Yasir A. Syed, Sung Ung Kang, Dieter Mitteregger, Raluca Vig, Robin J. M. Franklin, Friedrich Altmann, Gert Lubec, and Mark R. Kotter. 2009. “Myelin-Mediated Inhibition of Oligodendrocyte Precursor Differentiation Can Be Overcome by Pharmacological Modulation of Fyn-RhoA and Protein Kinase C Signalling.” Brain 132:445–81. doi:10.1093/brain/awn334.

5. Barres, B. A, I. K. Hart, H. S. R. Coles, J. F. Burne, J. T. Voyvodic, W. D. Richardson, and M. C. Raff. 1992. “Cell Death and Control of Cell Survival in the Oligodendrocyte Lineage.” Cell 70:31– 46.

6. Berghoff, Stefan A., Lena Spieth, Ting Sun, Leon Hosang, Lennart Schlaphoff, Constanze Depp, Tim Düking, Jan Winchenbach, Jonathan Neuber, David Ewers, Patricia Scholz, Franziska van der Meer, Ludovico Cantuti-Castelvetri, Andrew O. Sasmita, Martin Meschkat, Torben Ruhwedel, Wiebke Möbius, Roman Sankowski, Marco Prinz, Inge Huitinga, Michael W. Sereda, Francesca Odoardi, Till Ischebeck, Mikael Simons, Christine Stadelmann-Nessler, Julia M. Edgar, Klaus Armin Nave, and Gesine Saher. 2021. “Microglia Facilitate Repair of Demyelinated Lesions via Post-Squalene Sterol Synthesis.” Nature Neuroscience 24(1):47–60. doi:10.1038/S41593-020-00757-6.

7. Berghoff, Stefan, Lena Spieth, Ting Sun, Till Ischebeck, Julia M. Edgar, and Gesine Saher. 2021. “Neuronal Cholesterol Synthesis Is Essential for Repair of Chronically Demyelinated Lesions in Mice.” Cell Reports 37. doi:10.1016/j.celrep.2021.109889.

8. Blakemore, W. F., and R. J. M. Franklin. 2008. “Remyelination in Experimental Models of Toxin-Induced Demyelination.” Current Topics in Microbiology and Immunology 318:193–212. doi:10.1007/978-3-540-73677-6_8.

9. Bradl, Monika, and Hans Lassmann. 2010. “Oligodendrocytes: Biology and Pathology.” Acta Neuropathologica 119:37–53. doi:10.1007/s00401-009-0601-5.

10. Chia, Kelda, Anna Klingseisen, Dirk Sieger, and Josef Priller. 2022. “Zebrafish as a Model Organism for Neurodegenerative Disease.” Frontiers in Molecular Neuroscience 15. doi:10.3389/FNMOL.2022.940484/FULL.

11. Dimou, Leda, and Magdalena Götz. 2014. “Glial Cells as Progenitors and Stem Cells: New Roles in the Healthy and Diseased Brain.” Physiological Reviews 94(3):709–37. doi:10.1152/PHYSREV.00036.2013/ASSET/IMAGES/Z9J0021426900004.JPEG.

12. Dobin, Alexander, Carrie A. Davis, Felix Schlesinger, Jorg Drenkow, Chris Zaleski, Sonali Jha, Philippe Batut, Mark Chaisson, and Thomas R. Gingeras. 2012. “STAR: Ultrafast Universal RNA-Seq Aligner.” Bioinformatics 29(1):15. doi:10.1093/BIOINFORMATICS/BTS635.

13. Domercq, María, María Victoria Sánchez-Gómez, Catherine Sherwin, Estibaliz Etxebarria, Robert Fern, and Carlos Matute. 2007. “System Xc− and Glutamate Transporter Inhibition Mediates Microglial Toxicity to Oligodendrocytes.” The Journal of Immunology 178(10):6549–56. doi:10.4049/JIMMUNOL.178.10.6549.

14. Doszyn, O., T. Dulski, and J. Zmorzynska. 2024. “Diving into the Zebrafish Brain: Exploring Neuroscience Frontiers with Genetic Tools, Imaging Techniques, and Behavioral Insights.” Frontiers in Molecular Neuroscience 17. doi:10.3389/FNMOL.2024.1358844/FULL.

15. Ellett, Felix, Luke Pase, John W. Hayman, Alex Andrianopoulos, and Graham J. Lieschke. 2011. “Mpeg1 Promoter Transgenes Direct Macrophage-Lineage Expression in Zebrafish.” Blood 117(4). doi:10.1182/BLOOD-2010-10-314120.

16. Emery, Ben. 2010. “Regulation of Oligodendrocyte Differentiation and Myelination.” Science 330:779–82. doi:10.1126/science.1190927.

17. Falcão, Ana Mendanha, David van Bruggen, Sueli Marques, Mandy Meijer, Sarah Jäkel, Eneritz Agirre, Samudyata, Elisa M. Floriddia, Darya P. Vanichkina, Charles ffrench-Constant, Anna Williams, André Ortlieb Guerreiro-Cacais, and Gonçalo Castelo-Branco. 2018. “Disease-Specific Oligodendrocyte Lineage Cells Arise in Multiple Sclerosis.” Nature Medicine 24(12):1837–44. doi:10.1038/s41591-018-0236-y.

18. Faria-Pereira, Andreia, and Vanessa A. Morais. 2022. “Synapses: The Brain’s Energy-Demanding Sites.” International Journal of Molecular Sciences 23(7)(Research of Mitochondrial Function, Structure, Dynamics and Intracellular Organization):3627. 10.3390/ijms23073627.

19. Fernández-Castañeda, Anthony, Megan S. Chappell, Dorian A. Rosen, Scott M. Seki, Rebecca M. Beiter, David M. Johanson, Delaney Liskey, Emily Farber, Suna Onengut-Gumuscu, Christopher C. Overall, Jeffrey L. Dupree, and Alban Gaultier. 2020. “The Active Contribution of OPCs to Neuroinflammation Is Mediated by LRP1.” Acta Neuropathologica 139(2):365–82. doi:10.1007/S00401-019-02073-1/FIGURES/8.

20. Franklin, Robin J. M., and Charles Ffrench-Constant. 2017. “Regenerating CNS Myelin - From Mechanisms to Experimental Medicines.” Nature Reviews Neuroscience 18(12):753–69.

21. Fünfschilling, Ursula, Lotti M. Supplie, Don Mahad, Susann Boretius, Aiman S. Saab, Julia Edgar, Bastian G. Brinkmann, Celia M. Kassmann, Iva D. Tzvetanova, Wiebke Möbius, Francisca Diaz, Dies Meijer, Ueli Suter, Bernd Hamprecht, Michael W. Sereda, Carlos T. Moraes, Jens Frahm, Sandra Goebbels, and Klaus Armin Nave. 2012. “Glycolytic Oligodendrocytes Maintain Myelin and Long-Term Axonal Integrity.” Nature 2012 485:7399 485(7399):517–21. doi:10.1038/nature11007.

22. Gaultier, Alban, Xiaohua Wu, Natacha Le Moan, Shinako Takimoto, Gatambwa Mukandala, Katerina Akassoglou, W. Marie Campana, and Steven L. Gonias. 2009. “Low-Density Lipoprotein Receptor-Related Protein 1 Is an Essential Receptor for Myelin Phagocytosis.” Journal of Cell Science 122(8):1155–62. doi:10.1242/jcs.040717.

23. Gomez-Sanchez, Jose A., Lucy Carty, Marta Iruarrizaga-Lejarreta, Marta Palomo-Irigoyen, Marta Varela-Rey, Megan Griffith, Janina Hantke, Nuria Macias-Camara, Mikel Azkargorta, Igor Aurrekoetxea, Virginia Gutiérrez De Juan, Harold B. J. Jefferies, Patricia Aspichueta, Félix Elortza, Ana M. Aransay, María L. Martínez-Chantar, Frank Baas, José M. Mato, Rhona Mirsky, Ashwin Woodhoo, and Kristján R. Jessen. 2015. “Schwann Cell Autophagy, Myelinophagy, Initiates Myelin Clearance from Injured Nerves.” Journal of Cell Biology 210(1):153–68. doi:10.1083/jcb.201503019.

24. Hayashi, Teruo, and Tsung-Ping Su. 2004. “Sigma-1 Receptors at Galactosylceramide-Enriched Lipid Microdomains Regulate Oligodendrocyte Differentiation.” PNAS 101:14949–54.

25. Hayashi, Teruo, and Tsung Ping Su. 2007. “Sigma-1 Receptor Chaperones at the ER-Mitochondrion Interface Regulate Ca(2+) Signaling and Cell Survival.” Cell 131(3):596–610. doi:10.1016/J.CELL.2007.08.036.

26. Hofmann, Kristina, Rosalia Rodriguez-Rodriguez, Anne Gaebler, Núria Casals, Anja Scheller, and Lars Kuerschner. 2017. “Astrocytes and Oligodendrocytes in Grey and White Matter Regions of the Brain Metabolize Fatty Acids.” Nature Scientific Reports 7. doi:10.1038/s41598-017-11103-5.

27. Hornung, Veit, Franz Bauernfeind, Annett Halle, Eivind O. Samstad, Hajime Kono, Kenneth L. Rock, Katherine A. Fitzgerald, and Eicke Latz. 2008. “Silica Crystals and Aluminum Salts Activate the NALP3 Inflammasome through Phagosomal Destabilization.” Nature Immunology 9(8):847–56. doi:10.1038/ni.1631.

28. Hou, Jinchao, Yingyue Zhou, Zhangying Cai, Steven J. Van Dyken, Maxim N. Artyomov, Marco Colonna Correspondence, Marina Terekhova, Amanda Swain, Prabhakar S. Andhey, Rafaela M. Guimaraes, Alina Ulezko Antonova, Tian Qiu, Sanja Sviben, Gregory Strout, James A. J. Fitzpatrick, Yun Chen, Susan Gilfillan, Do-Hyun Kim, and Marco Colonna. 2023. “Transcriptomic Atlas and Interaction Networks of Brain Cells in Mouse CNS Demyelination and Remyelination.” doi:10.1016/j.celrep.2023.112293.

29. Jaillard, C., S. Harrison, B. Stankoff, M. S. Aigrot, A. R. Calver, G. Duddy, F. S. Walsh, M. N. Pangalos, N. Arimura, K. Kaibuchi, B. Zalc, and C. Lubetzki. 2005. “Cellular/Molecular Edg8/S1P5: An Oligodendroglial Receptor with Dual Function on Process Retraction and Cell Survival.” doi:10.1523/JNEUROSCI.4645-04.2005.

30. Ji, Xiao-Yu, Yu-Xin Guo, Li-Bin Wang, Wen-Cheng Wu, Jia-Qi Wang, Jin He, Rui Gao, Javad Rasouli, Meng-Yuan Gao, Zhen-Hai Wang, Dan Xiao, Wei-Feng Zhang, Bogoljub Ciric, Yuan Zhang, and Xing Li. 2024. “Microglia-Derived Exosomes Modulate Myelin Regeneration via MiR-615-5p/MYRF Axis.” Journal of Neuroinflammation 21–29. doi:10.1186/s12974-024-03019-5.

31. Kent, Sarah A., and Veronique E. Miron. 2024. “Microglia Regulation of Central Nervous System Myelin Health and Regeneration.” Nature Reviews Immunology 24(1):49–63.

32. Kimmel, Charles B., William W. Ballard, Seth R. Kimmel, Bonnie Ullmann, and Thomas F. Schilling. 1995. “Stages of Embryonic Development of the Zebrafish.” doi:10.1002/aja.1002030302.

33. Klosinski, Lauren P., Jia Yao, Fei Yin, Alfred N. Fonteh, Michael G. Harrington, Trace A. Christensen, Eugenia Trushina, and Roberta Diaz Brinton. 2015. “White Matter Lipids as a Ketogenic Fuel Supply in Aging Female Brain: Implications for Alzheimer’s Disease.” EBioMedicine 2(12):1888–1904. doi:10.1016/J.EBIOM.2015.11.002/ASSET/19C02D37-7800-4519-ADC4-ED79C8984A9B/MAIN.ASSETS/GR12.JPG.

34. Kotter, Mark R., Wen Wu Li, Chao Zhao, and Robin J. M. Franklin. 2006. “Myelin Impairs CNS Remyelination by Inhibiting Oligodendrocyte Precursor Cell Differentiation.” Journal of Neuroscience 26(1):328–32. doi:10.1523/JNEUROSCI.2615-05.2006.

35. Lassmann, Hans, and Monika Bradl. 2017. “Multiple Sclerosis: Experimental Models and Reality.” Acta Neuropathologica 133(2):223–44. doi:10.1007/S00401-016-1631-4.

36. Lee, Youngjin, Brett M. Morrison, Yun Li, Sylvain Lengacher, Mohamed H. Farah, Paul N. Hoffman, Yiting Liu, Akivaga Tsingalia, Lin Jin, Ping Wu Zhang, Luc Pellerin, Pierre J. Magistretti, and Jeffrey D. Rothstein. 2012. “Oligodendroglia Metabolically Support Axons and Contribute to Neurodegeneration.” Nature 487(7408):443–48. doi:10.1038/NATURE11314.

37. Liao, Yang, Gordon K. Smyth, and Wei Shi. 2014. “FeatureCounts: An Efficient General Purpose Program for Assigning Sequence Reads to Genomic Features.” Bioinformatics 30(7):923–30. doi:10.1093/BIOINFORMATICS/BTT656.

38. Liberzon, Arthur, Chet Birger, Helga Thorvaldsdóttir, Mahmoud Ghandi, Jill P. Mesirov, and Pablo Tamayo. 2015. “The Molecular Signatures Database (MSigDB) Hallmark Gene Set Collection.” Cell Systems 1(6):417. doi:10.1016/J.CELS.2015.12.004.

39. Lin, Jing Ping, Yevgeniya A. Mironova, Peter Shrager, and Roman J. Giger. 2017. “LRP1 Regulates Peroxisome Biogenesis and Cholesterol Homeostasis in Oligodendrocytes and Is Required for Proper CNS Myelin Development and Repair.” ELife 6. doi:10.7554/ELIFE.30498.

40. Love, Michael I., Wolfgang Huber, and Simon Anders. 2014. “Moderated Estimation of Fold Change and Dispersion for RNA-Seq Data with DESeq2.” Genome Biology 15(12):1–21. doi:10.1186/S13059-014-0550-8/FIGURES/9.

41. Marisca, Roberta, Tobias Hoche, Eneritz Agirre, Laura Jane Hoodless, Wenke Barkey, Franziska Auer, Gonçalo Castelo-Branco, and Tim Czopka. 2020. “Functionally Distinct Subgroups of Oligodendrocyte Precursor Cells Integrate Neural Activity and Execute Myelin Formation.” Nature Neuroscience 2020 23:3 23(3):363–74. doi:10.1038/s41593-019-0581-2.

42. Mccarthy, Ken D., and Jean De Vellis. 1980. “Preparation of Separate Astroglial and Oligodendroglial Cell Cultures from Rat Cerebral Tissue.” Journal Cell Biology 85:890–902.

43. Meffre, Delphine, Ghjacumu Shackleford, Mehdi Hichor, Victor Gorgievski, Eleni T. Tzavara, Amalia Trousson, Abdel M. Ghoumari, Cyrille Deboux, Brahim Nait Oumesmar, Philippe Liere, Michael Schumacher, Etienne-Emile Baulieu, Frédéric Charbonnier, Julien Grenier, and Charbel Massaad. 2015. “Liver X Receptors Alpha and Beta Promote Myelination and Remyelination in the Cerebellum.” 112(24):7587–92. doi:10.1073/pnas.1424951112.

44. Molina-Gonzalez, Irene, Rebecca K. Holloway, Zoeb Jiwaji, Owen Dando, Sarah A. Kent, Katie Emelianova, Amy F. Lloyd, Lindsey H. Forbes, Ayisha Mahmood, Thomas Skripuletz, Viktoria Gudi, James A. Febery, Jeffrey A. Johnson, Jill H. Fowler, Tanja Kuhlmann, Anna Williams, Siddharthan Chandran, Martin Stangel, Andrew J. M. Howden, Giles E. Hardingham, and Veronique E. Miron. 2023. “Astrocyte-Oligodendrocyte Interaction Regulates Central Nervous System Regeneration.” Nature Communications 14. doi:10.1038/s41467-023-39046-8.

45. Moyon, Sarah, Anne Laure Dubessy, Marie Stephane Aigrot, Matthew Trotter, Jeffrey K. Huang, Luce Dauphinot, Marie Claude Potier, Christophe Kerninon, Stephane Melik Parsadaniantz, Robin J. M. Franklin, and Catherine Lubetzki. 2015. “Demyelination Causes Adult CNS Progenitors to Revert to an Immature State and Express Immune Cues That Support Their Migration.” The Journal of Neuroscience : The Official Journal of the Society for Neuroscience 35(1):4–20. doi:10.1523/JNEUROSCI.0849-14.2015.

46. Nave, Klaus Armin, Ebrahim Asadollahi, and Andrew Sasmita. 2023. “Expanding the Function of Oligodendrocytes to Brain Energy Metabolism.” Current Opinion in Neurobiology 83. doi:10.1016/J.CONB.2023.102782.

47. Nave, Klaus Armin, and Hauke B. Werner. 2014. “Myelination of the Nervous System: Mechanisms and Functions.” Pp. 503–33 in Annual Review of Cell and Developmental Biology. Vol. 30. Annual Reviews Inc.

48. Nelissen, Katherine, Monique Mulder, Ilse Smets, Silke Timmermans, Karen Smeets, Marcel Ameloot, and Jerome J. A. Hendriks. 2011. “Liver X Receptors Regulate Cholesterol Homeostasis in Oligodendrocytes.” Journal of Neuroscience Research. doi:10.1002/jnr.22743.

49. Nogaroli, Luciana, A. E. Larra, M. Yuelling, A. E. Jameel, Dennis Ae, Karen Gorse Ae, Shawn G. Payne Ae, and Babette Fuss. 2009. “Lysophosphatidic Acid Can Support the Formation of Membranous Structures and an Increase in MBP MRNA Levels in Differentiating Oligodendrocytes.” Neurochem Res 34:182–93. doi:10.1007/s11064-008-9772-z.

50. Norton, W. T., and Shirley E. Poduslo. 1973. “Myelination in the Rat Brain: Method of Myelin Isolation.” Journal of Neurochemistry 21(4):749–57. doi:10.1111/J.1471-4159.1973.TB07519.X.

51. Nugent, A., Karin Lin, Bettina van Lengerich, Joseph W. Lewcock, Kathryn M. Monroe, Gilbert Di Paolo, Alicia A. Nugent, Steve Lianoglou, Laralynne Przybyla, Sonnet S. Davis, Ceyda Llapashtica, Junhua Wang, Do Jin Kim, Dan Xia, Anthony Lucas, Sulochanadevi Baskaran, Patrick CG Haddick, Melina Lenser, Timothy K. Earr, Ju Shi, Jason C. Dugas, Benjamin J. Andreone, Todd Logan, Hilda O. Solanoy, Hang Chen, Ankita Srivastava, Suresh B. Poda, Pascal E. Sanchez, Ryan J. Watts, Thomas Sandmann, and Giuseppe Astarita. 2020. “TREM2 Regulates Microglial Cholesterol Metabolism upon Chronic Phagocytic Challenge.” Neuron 105:837–54. doi:10.1016/j.neuron.2019.12.007.

52. Pandey, Shristi, Kimberle Shen, Seung-Hye Lee, Christopher J. Bohlen, Tracy J. Yuen, Brad A. Friedman Correspondence, Yun-An A. Shen, Yuanyuan Wang, Marcos Otero-García, Natalya Kotova, Stephen T. Vito, Benjamin I. Laufer, Dwight F. Newton, Mitchell G. Rezzonico, Jesse E. Hanson, Joshua S. Kaminker, and Brad A. Friedman. 2022. “Disease-Associated Oligodendrocyte Responses across Neurodegenerative Diseases.” Cell Reports 40. doi:10.1016/j.celrep.2022.111189.

53. Philips, Thomas, and Jeffrey D. Rothstein. 2017. “Oligodendroglia: Metabolic Supporters of Neurons.” Journal of Clinical Investigation 127(9):3271–80.

54. Plemel, Jason R., Sohrab B. Manesh, Joseph S. Sparling, and Wolfram Tetzlaff. 2013. “Myelin Inhibits Oligodendroglial Maturation and Regulates Oligodendrocytic Transcription Factor Expression.” GLIA 61(9):1471–87. doi:10.1002/glia.22535.

55. Prada, Ilaria, Ladan Amin, Roberto Furlan, Giuseppe Legname, Claudia Verderio, and Dan Cojoc. 2016. “A New Approach to Follow a Single Extracellular Vesicle - Cell Interaction Using Optical Tweezers.” BioTechniques 60(1):35–35. doi:10.2144/000114371.

56. Ralhan, Isha, Chi-Lun Chang, Jennifer Lippincott-Schwartz, and Maria S. Ioannou. 2021. “Lipid Droplets in the Nervous System.” doi:10.1083/jcb.202102136.

57. Ramos-Cabrer, Pedro, Alberto Cabrera-Zubizarreta, Daniel Padro, Mario Matute-González, Alfredo Rodríguez-Antigüedad, and Carlos Matute. 2025. “Reversible Reduction in Brain Myelin Content upon Marathon Running.” Nature Metabolism. doi:10.1038/s42255-025-01244-7.

58. Rao, Vijayaraghava T. S., Damla Khan, Qiao Ling Cui, Shih Chieh Fuh, Shireen Hossain, Guillermina Almazan, Gerhard Multhaup, Luke M. Healy, Timothy E. Kennedy, and Jack P. Antel. 2017. “Distinct Age and Differentiation-State Dependent Metabolic Profiles of Oligodendrocytes under Optimal and Stress Conditions.” PloS One 12(8). doi:10.1371/JOURNAL.PONE.0182372.

59. Rinholm, Johanne E., Nicola B. Hamilton, Nicoletta Kessaris, William D. Richardson, Linda H. Bergersen, and David Attwell. 2011. “Regulation of Oligodendrocyte Development and Myelination by Glucose and Lactate.” doi:10.1523/JNEUROSCI.3516-10.2011.

60. Sánchez-Gómez, Maria Victoria, Mari Paz Serrano, Elena Alberdi, Fernando Pérez-Cerdá, and Carlos Matute. 2018. “Isolation, Expansion, and Maturation of Oligodendrocyte Lineage Cells Obtained from Rat Neonatal Brain and Optic Nerve.” Pp. 95–113 in Myelin. Methods in Molecular Biology. Vol. 1791, Methods in Molecular Biology, edited by A. Woodhoo. New York, NY: Humana Press.

61. Simons, Mikael, and Klaus Armin Nave. 2016. “Oligodendrocytes: Myelination and Axonal Support.” Cold Spring Harbor Perspectives in Biology 8(1).

62. Stadelmann, C., S. Timmler, A. Barrantes-Freer, and M. Simons. 2019. “Myelin in the Central Nervous System: Structure, Function, and Pathology.” Physiol Rev 99:1381–1431. doi:10.1152/physrev.00031.2018.-Oligodendro.

63. Tang, Min, Xin Dong, Lanbo Xiao, Zheqiong Tan, Xiangjian Luo, Lifang Yang, Wei Li, Feng Shi, Yueshuo Li, Lin Zhao, Na Liu, Qianqian Du, Longlong Xie, Jianmin Hu, Xinxian Weng, Jia Fan, Jian Zhou, Qiang Gao, Weizhong Wu, Xin Zhang, Weihua Liao, Ann M. Bode, and Ya Cao. 2022. “CPT1A-Mediated Fatty Acid Oxidation Promotes Cell Proliferation via Nucleoside Metabolism in Nasopharyngeal Carcinoma.” Cell Death & Disease 13(4):331. doi:10.1038/S41419-022-04730-Y.

64. Tekkök, Selva Baltan, Angus M. Brown, Ruth Westenbroek, Luc Pellerin, and Bruce R. Ransom. 2005. “Transfer of Glycogen-Derived Lactate from Astrocytes to Axons via Specific Monocarboxylate Transporters Supports Mouse Optic Nerve Activity.” Journal of Neuroscience Research 81(5):644–52. doi:10.1002/JNR.20573.

65. Tepavčevic, Vanja. 2021. “Oligodendroglial Energy Metabolism and (Re)Myelination.” doi:10.3390/life.

66. Terada, Nobuo, Karen Baracskay, Mike Kinter, Shona Melrose, Peter J. Brophy, Claude Boucheix, Carl Bjartmar, Grahame Kidd, and Bruce D. Trapp. 2002. “The Tetraspanin Protein, CD9, Is Expressed by Progenitor Cells Committed to Oligodendrogenesis and Is Linked to Β1 Integrin, CD81, and Tspan-2.” GLIA 40(3):350–59. doi:10.1002/GLIA.10134.

67. Torre-Fuentes, L., L. Moreno-Jiménez, V. Pytel, J. A. Matías-Guiu, U. Gómez-Pinedo, and J. Matías-Guiu. 2020. “Experimental Models of Demyelination and Remyelination.” Neurología (English Edition) 35(1):32–39. doi:10.1016/J.NRLENG.2019.03.007.

68. Trivedi, Purvi C., Jordan J. Bartlett, and Thomas Pulinilkunnil. 2020. “Lysosomal Biology and Function: Modern View of Cellular Debris Bin.” Cells 9:1131. doi:10.3390/cells9051131.

69. Vagionitis, Stavros, and Tim Czopka. 2018. “Visualization and Time-Lapse Microscopy of Myelinating Glia in Vivo in Zebrafish.” Methods in Molecular Biology 1791:25–35. doi:10.1007/978-1-4939-7862-5_3/FIGURES/2.

70. Young, Kaylene M., Konstantina Psachoulia, Richa B. Tripathi, Sara-Jane Dunn, Lee Cossell, David Attwell, Koujiro Tohyama, and William D. Richardson. 2013. “Oligodendrocyte Dynamics in the Healthy Adult CNS: Evidence for Myelin Remodeling.” Neuron 77:873–85. doi:10.1016/j.neuron.2013.01.006.

71. Yu, Fang, Yangfan Wang, Anne R. Stetler, Rehana K. Leak, Xiaoming Hu, and Jun Chen. 2022. “Phagocytic Microglia and Macrophages in Brain Injury and Repair.” CNS Neurosci Ther 28:1279–93. doi:10.1111/cns.13899.

72. Yung, Yun C., Nicole C. Stoddard, Hope Mirendil, and Jerold Chun. 2015. “Lysophosphatidic Acid (LPA) Signaling in the Nervous System.” Neuron 85(4):669. doi:10.1016/J.NEURON.2015.01.009.

73. Zeis, Thomas, Lukas Enz, and Nicole Schaeren-Wiemers. 2015. “The Immunomodulatory Oligodendrocyte.” Brain Research 139–48. doi:10.1016/j.brainres.2015.09.021.

74. Zirngibl, Martin, Peggy Assinck, Anastasia Sizov, Andrew V Caprariello, and Jason R. Plemel. 2021. “Oligodendrocyte Death and Myelin Loss in the Cuprizone Model: An Updated Overview of the Intrinsic and Extrinsic Causes of Cuprizone Demyelination.” doi:10.1186/s13024-022-00538-8.

