## Supplementary figures for "Internalization of exogenous myelin by oligodendroglia promotes lineage progression"

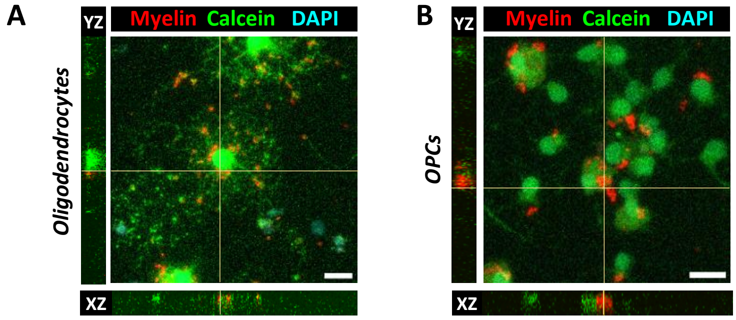


***Supplementary Figure 1. Internalization of exogenous myelin by oligodendroglial cells.*** (**A–B**) Orthogonal confocal views showing exogenous myelin internalized within oligodendrocytes (**A**) and OPCs (**B**), visualized using Calcein-AM to label live cells. Scale bar = 20 µm.

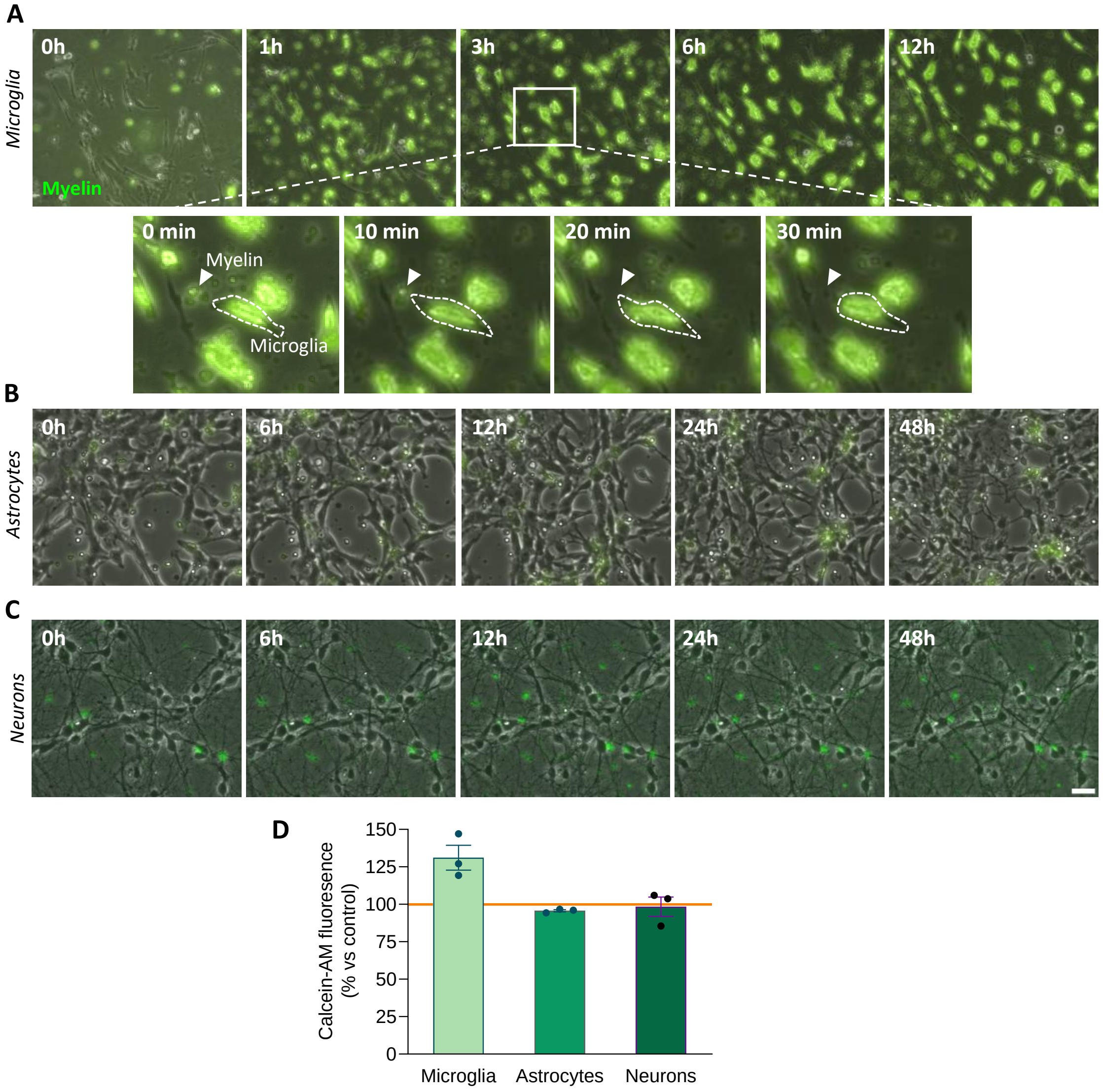


**Supplementary Figure 2. Internalization of exogenous myelin by glial cells and neurons**. (**A–C**) Time-lapse imaging showing the internalization of myelin debris by microglia (A), astrocytes (B), and neurons (**C**). (**D**) Cell viability assay performed on each cell type following 48 hours of exposure to myelin debris. Scale bar = 5 µm.

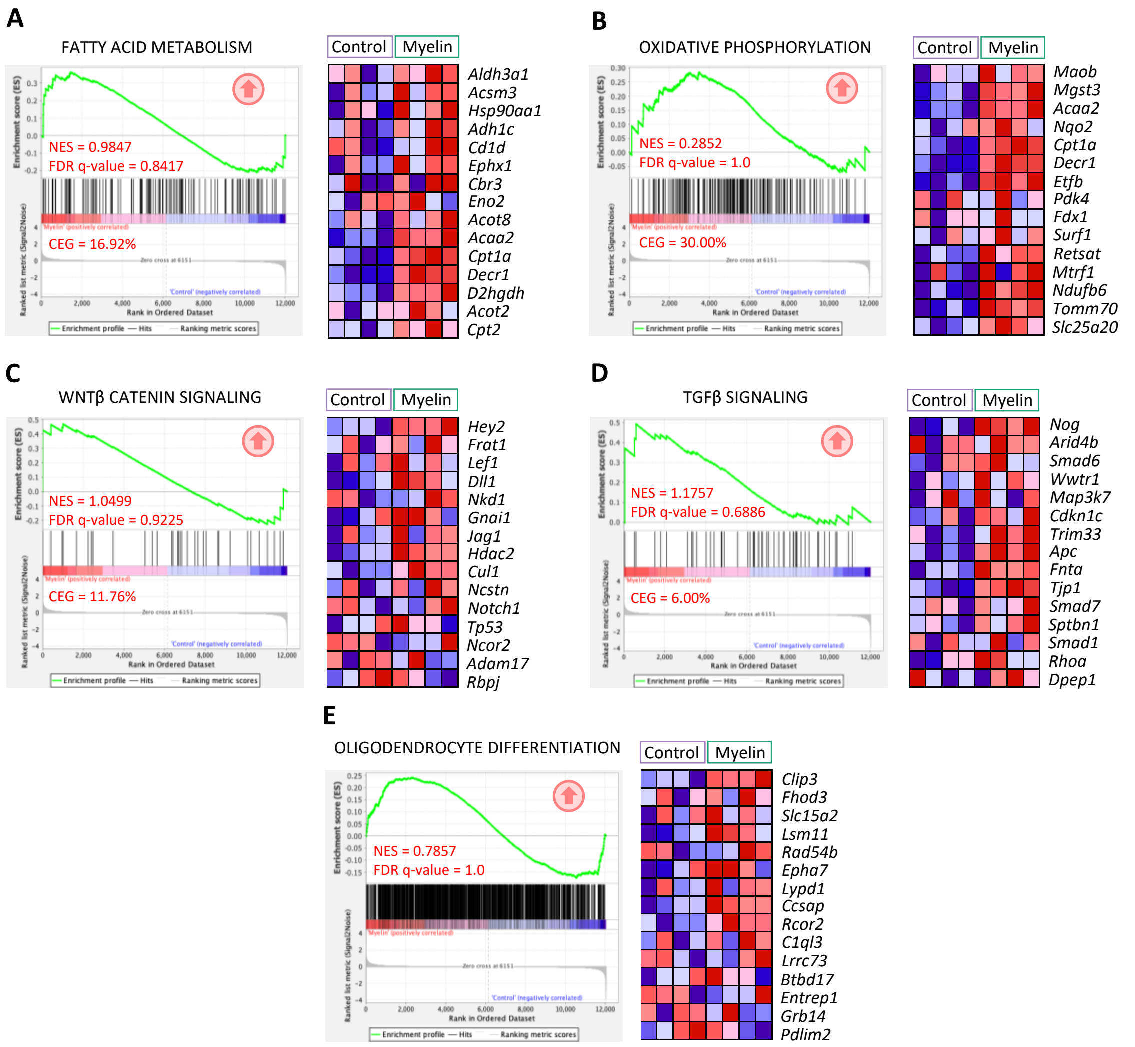


**Supplementary Figure 3. Metabolic and differentiation transcriptional profile of oligodendrocytes after myelin exposure.** Enrichment plots (left) for selected Hallmark gene sets (**A-D**) or specific signatures (**E**) with corresponding NES, FDR q-values, and Core Enrichment Genes (CEG). Heatmaps (right) show the top 15 most enriched genes for each pathway

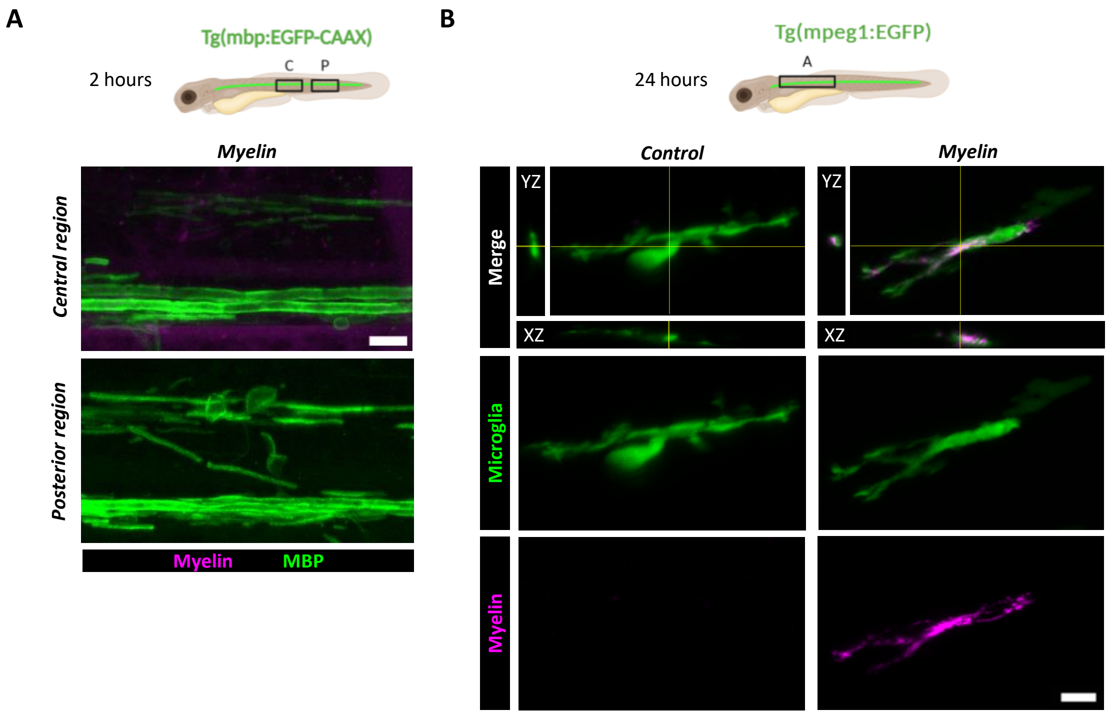


**Supplementary Figure 3. Internalization of exogenous myelin in the zebrafish model**. (**A**) Confocal imaging of the spinal cord 2 hours after cerebroventricular injection reveals the absence of fluorescent myelin in the central and posterior regions of the animal. (**B**) Microglia internalizing myelin debris 24 hours post-injection in the spinal cord of *Tg(mpeg1:EGFP)* zebrafish, which express EGFP in microglial cells. Scale bar = 10 µm
